# Human Neocortical Glutamatergic Neurons Revealed Through Multimodal Profiling

**DOI:** 10.64898/2026.01.08.698257

**Authors:** Rachel Dalley, Sarah Walling-Bell, Jeremy A. Miller, Femke Waleboer, Julian Thijssen, Audrey McCutcheon, Emily Gelfand, Matt Mallory, Cristina Radaelli, Xiao-Ping Liu, Scott Sawchuk, Lauren Alfiler, Julia Andrade, Jeanelle Ariza, Angela Ayala, Katherine Baker, Eliza Barkan, Stuard Barta, Kyla Berry, Darren Bertagnolli, Kris Bickley, Madeline Bixby, Krista Blake, Jasmine Bomben, Krissy Brouner, Agata Budzillo, Jazmin Campos, Trangthanh Cardenas, Daniel Carey, Tamara Casper, Anish Bhaswanth Chakka, Michael Clark, Tom V. Coopmans, Kirsten Crichton, Nick Dee, Nadezhda I. Dotson, Stan L.W. Driessens, Tom Egdorf, Lauren Ellingwood, Chris English, Rachel Enstrom, Rebecca de Frates, Anna A. Galakhova, Amanda Gary, Jessica Gloe, Jeff Goldy, Hong Gu, Kristen Hadley, Samantha D. Hastings, Tim S. Heistek, Dijon Hill, Windy Ho, Alvin Huang, Daan C. Jol, Atlas Jordan, Zoe C. Juneau, Matthew Jungert, Madhav Kannan, Sara Kebede, Ayoub J. Khalil, Lisa Kim, Mean-Hwan Kim, Megan Koch, Ágnes Katalin Kocsis, Matthew Kroll, Aditi Kumar, Gabriela Leon, Jocelin Malone, Rusty Mann, Michelle Maxwell, Medea McGraw, Delissa A. McMillen, Eline J. Mertens, Verjinia D. Metodieva, Miranda R. Moore, Alice Mukora, Kamiliam Nasirova, Lindsay Ng, Aaron Oldre, Alana Oyama, Daniel Park, Elliot Phillips, Christina Alice Pom, Lydia Potekhina, Ramkumar Rajanbabu, Shea T. Ransford, Ingrid Redford, Christine Rimorin, Dana Rocha, Augustin Ruiz, David Sandman, Natia Shamugia, Ana Rios Sigler, Josef Sulc, Tjerk P. Swinkels, Michael Tieu, Éva Toth, Alex Tran, Jessica Trinh, Herman Tung, Marshall VanNess, Sara Vargas, Maria Camila Vergara, Katherine Wadhwani, Grace Williams, Laurens Witter, Anubhav G. Amin, Charles Cobbs, Richard Ellenbogen, Samuel Emerson, Manuel Ferreira, Nathan W. Gouwens, Benjamin Grannan, Ryder P. Gwinn, Jason S. Hauptman, Sander Idema, Andrew L. Ko, Lauren Kruse, Brian Long, David Noske, Jeffrey G. Ojemann, Anoop Patel, Jacob J. Ruzevick, Dan L. Silbergeld, Kimberly A. Smith, Niels Verburg, Jack Waters, Tim Jarsky, Boudewijn Lelieveldt, Gábor Tamás, Huibert D. Mansvelder, Natalia A. Goriounova, Christiaan P.J. de Kock, Nikolai C. Dembrow, Jonathan T. Ting, Hongkui Zeng, Ed S. Lein, Brian Kalmbach, Staci A. Sorensen, Brian R. Lee

## Abstract

The human neocortex underlies higher cognition and is the engine of complex thought. Yet our understanding of its neuronal diversity is limited by sparse access to tissue, inconsistent sampling across studies, and a lack of multiple modality data. Although single-cell transcriptomic taxonomies are an important framework for characterizing cell type diversity, transcriptomic information alone cannot reveal the cellular properties that define neuronal computations. To address this, we performed Patch-seq, a method for collecting Morphology, Electrophysiology, and Transcriptomic data from a single neuron. We focused on glutamatergic, neocortical, excitatory neurons, the principal long-range projecting neurons of the cortex, and systematically integrated their morphoelectric features with transcriptomic identity. In combination with spatial transcriptomic data, we interrogated 39 of 42 transcriptomically-defined neuron types with a layer-centric perspective. Morphoelectric properties, such as cortical depth, apical dendrite structure, and excitability clearly distinguish transcriptomic subclasses and support many finer transcriptomic types. Morphoelectric properties are influenced by spatial location in supragranular layers, while deeper layers exhibit greater heterogeneity. Cross-species comparisons reveal conserved subclass organization but pronounced differences in apical dendrite arborization between mouse and human, and surprising similarities between human and macaque. Together, these datasets provide a unified multimodal reference that advances our understanding of human cortical circuitry and establishes a foundation for experimental and computational studies of human brain function and disease.

## Introduction

The human neocortex, organized into six layers, is responsible for complex cognitive functions. Elucidating the cell type identity and functional properties of the neurons that are fundamental to those functions is essential for understanding cortical computation and for translating those insights to human health and behavior.

Though there have been several studies investigating transcriptomically-defined neuronal cell types in individual layers of human cortex^1–4^, much of what we know about the phenotypic properties of these cell types, such as their distribution, shape, and physiology, has come from rodent studies^5–8^. A comprehensive characterization of multimodal neuronal types is needed for human neocortex to accelerate our understanding of human cortical circuit function, species differences, and disease mechanisms.

Progress in understanding human neurons has been challenging due in large part to limited tissue availability. There have also been insufficient methods for targeting specific cell types, though this is changing rapidly^9–11^. Also, neuronal types have historically been defined by individual features, such as one or a few marker genes, laminar position, major axonal projection type, and/or electrophysiological behavior, making cross-study comparisons difficult^12,13^. Single-cell transcriptomics has transformed our ability to identify neuronal cell types and connect them across species and developmental time points^14–22^. However, transcriptomic data alone cannot reveal the cellular properties, such as morphology and electrophysiology, that facilitate an understanding of input–output transformations and how neurons embed within a network. Linking morpho-electric properties to transcriptomic identity yields a more complete picture of the cell type landscape.

In this study we focused on glutamatergic excitatory neurons, the neocortex’s primary source of long-distance, inter-areal communication. Traditionally, these neurons have been identified in rodent studies by their axon projections ^18,23,24^, with human assignments recently inferred from transcriptomic homology^15,16^. Our prior work on Layer 2/3 (L2/3) and L5 human glutamatergic neurons identified strong correspondence between morphological, electrophysiological and/or transcriptomic features^1,4^. We also found that morpho-electric properties of human L5 Extratelencephalic (ET) neurons were conserved in rodents, and that L5 cell type differences were often stronger within species than between them^4^. These findings offer a foundation for the present analysis, which provides a more comprehensive catalog of glutamatergic cortical neurons across all layers, mapped to a high-resolution transcriptomic taxonomy with more refined neuronal types and alignment with multimodal cell types in mouse and macaque.

For this work, we used Patch-seq^25,26^ on neurosurgically resected human tissue. We collected Morphology (M), Electrophysiology (E) and Transcriptomics (T) data, collectively referred to as MET, from the same neuron. Building on our previous morphoelectric characterization^1,4^ we systematically integrate morphoelectric properties with transcriptomic identity layer-by-layer for 39 of the 42 transcriptomically-defined excitatory neuron types described for human middle temporal gyrus (MTG). We identify key functional properties that distinguish L2 from L3 Intratelencephalic (IT) transcriptomic types. We characterize unique supertypes within the L4 IT subclass and highlight distinctive properties between L5 IT and L5/6 Near Projecting (NP) neurons. We evaluated non-traditional pyramidal morphologies and the diverse electrophysiological features of transcriptomically-defined L6 subclasses and find that electrophysiological features reliably predicted subclass identity. Cross-species comparisons reveal differences in the proportion of neurons with apical dendritic processes reaching L1 and underappreciated morphological similarities between human and macaque neurons. Finally, we include our dataset in Cytosplore, a tool that allows researchers to explore the dataset, examine relationships between all three modalities, and build upon our cell type definitions to answer questions toward advancing human health.

## Results

### Patch-seq profiling in human cortex

To identify morpho-electric signatures of transcriptomic cell types, we combined newly generated and previously published Patch-seq data^1,4^ from resected surgical tissue (n=1491 total; n=1125 unpublished; Figures 1A and 1B, Table S1). Neurons were mapped to a previously published human MTG Seattle Alzheimer’s Disease Brain Cell Atlas (SEA-AD) reference taxonomy^27^ that is organized into nested levels (broad class, intermediate subclass, and fine grained supertypes) using a hierarchical mapping approach (see Methods). After applying quality control gates (Figures S1A-S1D; Methods), our dataset consisted of transcriptomic, electrophysiology, and morphology modalities for 39, 37 and 36 of the 42 possible glutamatergic supertypes, respectively (Figure 1C). Neurons were predominantly sampled from temporal (71%) and frontal (16%) lobes (Figures 1D and S1E). Only neurons with intact apical dendrites were selected for dendritic reconstruction, facilitating comprehensive evaluation of dendritic morphology across supertypes (Figure S2). Most neurons (87%) were collected from acute ex vivo brain slices; to enhance sampling of glutamatergic types and increase tissue viability, an ex vivo brain slice culture paradigm was also used^11^. Consistent with previous results, electrophysiological features shifted subtly in culture. However, differences between preparations generally did not exceed differences between subclasses (Figure S3, Table S2)^2,11^, therefore electrophysiological analyses were aggregated across paradigms. Similarly, morphological features did not vary between preparations (Table S2).

**Figure 1.**
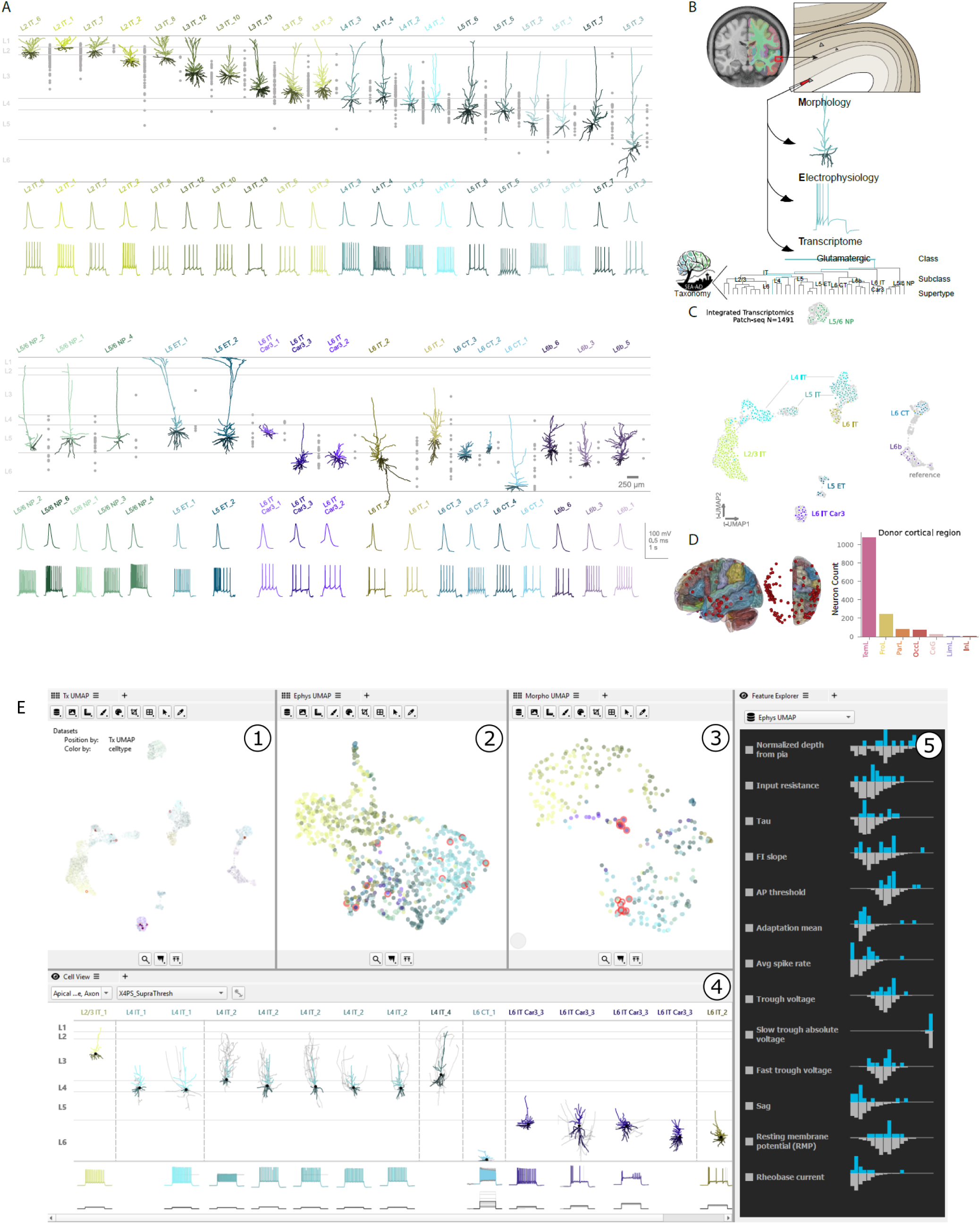
Patch-seq profiling in human cortex. A. from normalized depth from pia, while neurons with reconstructions are placed more precisely using layer alignment. Morphologies and action potential firing patterns arranged by subclass and ordered by their mean spatial transcriptomic laminar isodepth. B. Overview of the Patch-seq workflow. Transcriptomes from Patch-seq neurons were mapped to the SEA-AD reference taxonomy using a hierarchical mapping approach that iteratively and probabilistically assigns cell types to each taxonomy level using differentially expressed genes. C. The integrated UMAP, generated using the default Seurat (v4.4) workflow for data integration, of reference transcriptomic data (gray circles) and Patch-seq transcriptomes (colored by subclass). D. Human Common Coordinate Framework (CCF) showing centroids of donor tissue blocks and the numbers of neurons per cortical lobe used in this dataset. E. **Cytosplore Viewer:** Overview of Patch-seq data within Cytosplore Viewer. The software interface is organized into several linked sub-visualizations. 1. Integrated UMAP of reference and Patch-seq transcriptomes (colored by supertype). 2. UMAP of electrophysiological features for the subset of cells with electrophysiological recordings. 3. UMAP of morphological features for the subset of cells with morphological reconstructions. 4. Visualization of morphologies within an average cortical template (colored by supertype) and a corresponding representative voltage trace of the selected cells, when available. Different stimulus protocols can be selected to display their associated voltage responses and apical dendrite, basal dendrite and axon can be toggled for visualization. 5. Ranked list of either morphological or electrophysiological features, ordered from most to least salient for the selected cells. For each feature, the blue histogram shows the distribution of values for the selected cells, while the grey histogram shows the distribution across the entire dataset.

To enable interactive exploration and analysis of these rich, multimodal datasets, we provided the data as a Patch-seq explorer module within the Cytosplore Viewer visualization software (available at https://viewer.cytosplore.org)^28,29^(Figure 1E). The dataset can be viewed and analyzed within the Cytosplore Viewer or downloaded from public data repositories.

### Morphoelectric properties of transcriptomic subclasses

We first assessed glutamatergic neurons at the subclass level to capture broad differences in circuit organization and establish alignment with traditional groupings^23^. We examined spatial organization with MERFISH data and analyzed morphoelectric properties across nine transcriptomically-defined subclasses: L2/3 IT, L4 IT, L5 IT, L5/6 NP, L5 ET, L6 IT Car3, L6 IT, L6 Corticothalamic (CT) and L6b. These subclasses are conserved across species^15,16^ and the names, along with laminar position, were inferred from transcriptomic homology to mouse (Figure 2A). Human Patch-seq soma locations agreed with their subclass assignments (Figure 2B), though they also extended to adjacent layers, consistent with prior reports^1,16^.

**Figure 2.**
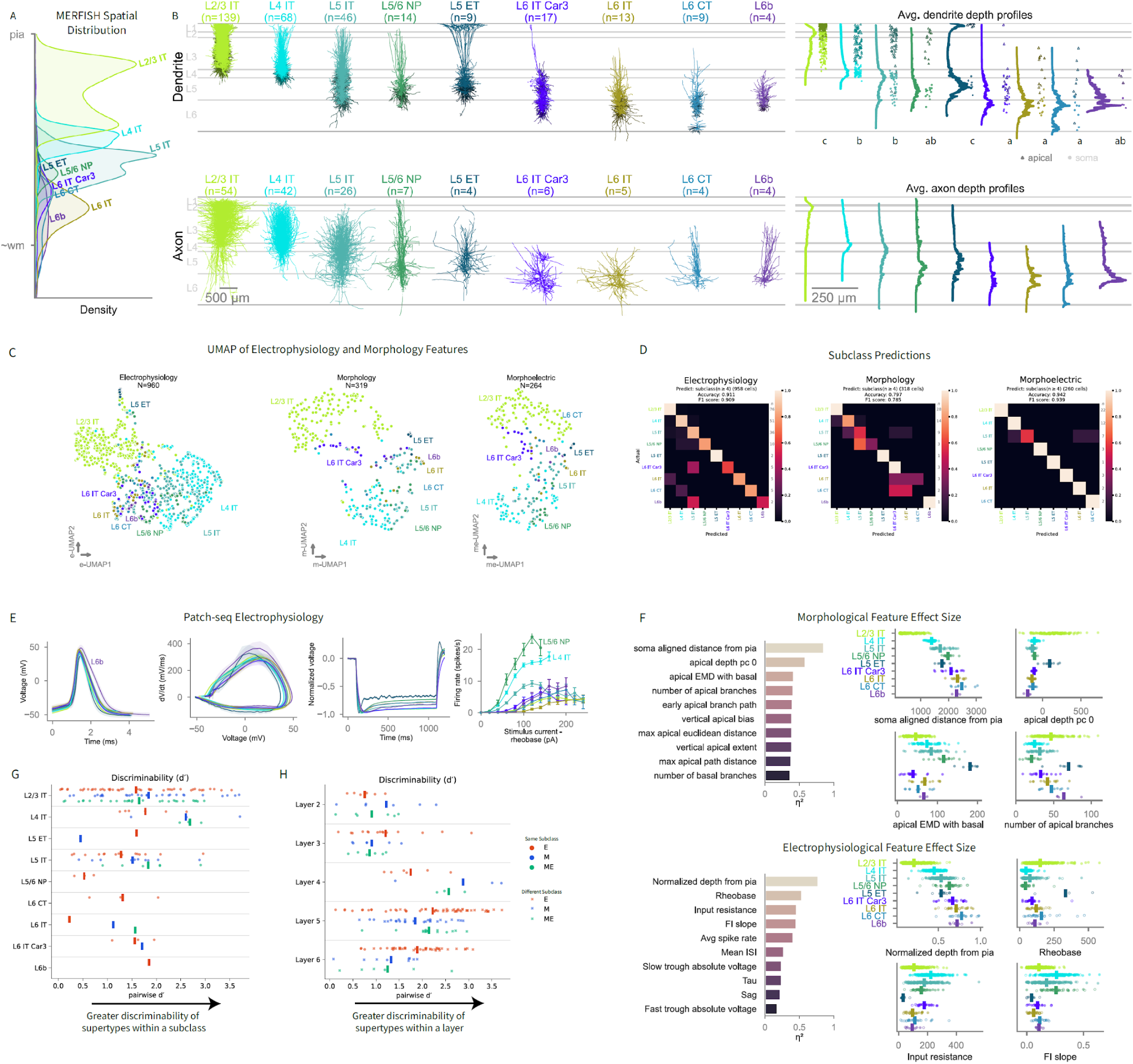
Morphoelectric properties of transcriptomic subclasses. A. Spatial transcriptomics density plot (kernel density estimate), colored by subclass. B. All reconstruction overlayed in the average layer template, dendrites (top) and local axons (bottom) by subclass. At right, apical dendrite tips nearest to layer 1 (triangles) and somas (circles) are shown, alongside histograms of average neurite length. A Kruskal-Wallis test p= 3.2 × 10⁻⁴³, FDR-corrected post hoc Dunn confirmed distinct laminar distributions of the apical tip nearest L1 across subclasses, with compact letter display groupings shown below, subclasses sharing the same letter are not significantly different. C. UMAP representations of electrophysiological (E), morphological (M), and combined morphoelectric (ME) feature spaces, with depth from pia included as a feature in each modality. D. Confusion matrix of a random forest classifier predicting subclass from electrophysiological, morphological and combined morphoelectric features. Only subclasses with at least four cells were included. Models were trained and evaluated on n = 958, 318, and 260 cells, achieving accuracies of 0.911, 0.797, and 0.942 and weighted F1 scores of 0.909, 0.785, and 0.939 for electrophysiological, morphological, and morphoelectric feature sets, respectively. E. Subclass action potentials with corresponding phase plot; voltage response to a 1-s hyperpolarizing current stimulus and the AP firing rate with increasing stimulus intensity above rheobase (f-I) curve averaged by their transcriptomically defined subclass. F. Feature importance analysis of morphological and electrophysiological features that best distinguish subclasses. A nonparametric ANOVA (Kruskal-Wallis H test) was used to calculate effect size (η²). Features are ranked by η², and the 10 largest values are shown at left. The top four features are visualized as strip plots at right with culture denoted as open circles and acute as closed circles (electrophysiology only). G. Pairwise supertype comparisons using the d′ discriminability metric, where higher values indicate greater separation. Pairwise d′ values are averaged to yield a mean d′ for each subclass shown as bars. H. Pairwise supertype comparisons using the d′ discriminability metric, where higher values indicate greater separation. Pairwise d′ values are averaged to yield a mean d′ for each layer shown as bars.

The laminar distribution of dendritic and axonal branches provides context for a neuron’s connectivity. The morphologies of human glutamatergic cortical neurons, aggregated by subclass, had distinct branching patterns. Each subclass had characteristic differences in dendrite and axon extent and arbor compactness (Figure 2B). Across subclasses, neurons differed significantly in the laminar position of their apical dendrite tip nearest to L1, with L2/3 IT and L5 ET neurons extending more superficially than all other subclasses. These data suggest that L2/3 IT and L5 ET subclasses receive more L1 inputs and are thus functionally distinct from other subclasses (Figure 2B, right). Local axonal distributions also offer insight into proximal connections and may offer clues about a neuron’s long-range innervation patterns^30^. To characterize local axons of glutamatergic subclasses, we reconstructed neurons with local axon greater than 2.2 mm in total length and found that local axon distribution differed across subclasses. For example, axons of L6 IT and L6 IT Car3 neurons elaborated horizontally, spanning >1mm, whereas axons of L5/6 NP, L6 CT and L6b neurons elaborated vertically, spanning multiple layers (Figures 2B and S2).

To visually assess the electrophysiological, morphological and combined morphoelectric landscape relative to transcriptomically-defined subclass, we used a Uniform Manifold Approximation and Projection (UMAP, Figure 2C). L2/3 IT, L4 IT and L5 ET subclasses occupied separate regions while the remaining infragranular subclasses were more intermingled, especially in electrophysiology space. Similar results were observed when soma cortical depth was excluded (Figure S4).

To evaluate the extent to which transcriptomic subclass can be predicted based on morphoelectric (ME) properties, we trained a random forest classifier with M, E and ME features. Subclasses with n<4 for each data modality were excluded. The combination of ME features (N = 260) provided the best prediction, with 94% accuracy (F1=.94). When trained exclusively on E features (N=958), the classifier achieved 91% accuracy (F1=.90); training exclusively on M features (N=318) resulted in 80% accuracy (F1=.78) (Figure 2D). Overall, transcriptomic subclass could be well-predicted based on morphoelectric properties. When misclassification occurred for morphology, it was often within the same axonal projection type and adjacent layer, for example, between L4 and L5 IT and within L6 subclasses.

Next, we compiled and averaged several electrophysiological features within their transcriptomically defined subclass for comparative analysis. Averages and standard deviation of single action potentials and corresponding phase plots revealed that average L6b action potentials (APs) had the greatest spike height and maximum rate of rise, but also the longest half widths (Figure 2E). Normalized voltage response to a 1-s hyperpolarizing current stimulus revealed substantial voltage sag in L5 ET neurons as has been shown previously^4,15^. Combined, these data underscored the finding that there were several distinctive electrophysiological properties that varied across glutamatergic subclasses.

To quantify which morphological and electrophysiological features best distinguished subclasses, we calculated effect size using eta squared (η²). We found that cortical soma depth explained the most variance between subclasses for both modalities. Features of the apical dendrites were particularly informative while basal dendrite features played a smaller role. Electrophysiology features related to neuronal excitability, such as the rheobase (the current required to initiate AP firing), Frequency-Current (F-I) slope and the average spike rate were also distinguishing features (Figure 2F). Rheobase exhibited the greatest heterogeneity across subclasses, with L4 IT and L5/6 NP requiring the least amount of current to elicit action potentials, while L2/3 IT and L5 ET required approximately three- to four-fold more current. Among the IT subclasses, we also observed heterogeneity in the intrinsic firing patterns. Analysis of suprathreshold properties revealed that L4 IT and L5/6 NP subclasses exhibited the fastest (AP) firing rates, with L5/6 NP also exhibiting spike-frequency accommodation, characterized by a gradual slowing of firing rate over time. L6 IT had the slowest firing rates, only exhibiting a burst at the onset of AP firing. Additionally, we noted heterogeneity in bursting within the L2/3 IT and L4 IT subclasses (Figure 1A). In addition to the firing properties of the neurons, passive electrophysiological properties such as input resistance and tau also distinguished subclasses, reflecting differences in neuronal size and ion channel distribution/density between them.

Given the strong effect of soma depth in distinguishing among subclasses, we reran classifiers without this feature to assess its importance. Classification accuracy for E and M decreased slightly, while ME stayed the same (Figure S4B). The same trend was observed when the classifiers were trained on a matched set of neurons for each modality (Figure S4C). The stability in the ME classifier reflects the fact that cortical depth is also correlated with many morphological and electrophysiological features.

Together, these results demonstrated that morphoelectric properties – particularly cortical depth, apical dendrite shape, and neuronal excitability – best distinguished subclass identity. Given the distribution of data points within each subclass, we sought to examine feature heterogeneity at the next, finer level of the transcriptomic hierarchy, that of supertypes.

### Morphoelectric properties of transcriptomic supertypes

Based on gene expression, neurons within each subclass can be further divided into transcriptomically-defined supertypes. How morphoelectric properties of human glutamatergic cortical neurons align to supertypes across cortical layers is unknown. To investigate this, we measured the distinctness of supertypes using a pairwise discriminability (d’) metric^31^, where higher values indicate greater separation in feature space. Within each subclass we compared every pair of supertypes and calculated d’, then averaged these values to get a mean d’. Using electrophysiological features alone, supertypes within L5/6 NP and L6 IT were the least distinct, while L4 IT and L6b supertypes were the most distinct. Morphology features similarly highlighted L4 IT supertypes as the most distinguishable (Figure 2G). Within L2/3 supertypes, we observed a wide range of d’ values indicating that some supertypes were well distinguished by morphoelectric features while others were not. To see if the wide range of L2/3 discriminability was unique or present in other layers, we also measured d’ for these supertypes within their primary layer (Figure 2H). In upper layers, L2 and L3 supertypes had lower discriminability. Supertypes within L4 were again the most discriminable using morphology and morphoelectric features. L5 supertypes had the highest level of discernability and the greatest range based on electrophysiology, reflecting the subclass diversity in infragranular layers. Together, these results highlight substantial subclass- and layer-dependent differences in how morphoelectric features correspond to supertypes, underscoring the need for closer examination of the 42 glutamatergic supertypes within the context of their cortical layer.

### Morphoelectric properties of supertypes in L2

The L2/3 IT subclass comprised the largest number of supertypes, ten compared to five defined in a previous human study^1^ and demonstrated the broadest range of discriminability among subclasses (Figure 2G). In humans, the supragranular layers are significantly expanded compared to mice, and variability in morphological, electrophysiological, and transcriptomic features align with the radial axis of L2/3^1^. We investigated whether L2/3 IT supertypes had distinct laminar and morphoelectric characteristics by analyzing over 650 Patch-seq neurons together with MERFISH spatial transcriptomic data.

A clear bimodal distribution of the L2/3 IT subclass in the spatial data prompted us to ask whether this pattern reflects contributions from specific supertypes (Figure 2A). When we ordered the ten supertypes by their mean isodepth, a value approximating cortical depth (methods), three supertypes, L2/3 IT_1, 6 and 7, were largely found in L2 (Figure 3A) and clustered together in the integrated transcriptomic UMAP (Figure 3B). Distinct L2 IT types have also been described for mouse^32^ and these human types exhibited similar morphological properties, suggesting that they are conserved across species (Figure S5A). These findings provided justification for treating these neurons as “L2” supertypes. Therefore, we refer to them as L2 IT_1, L2 IT_6 and L2 IT_7.

**Figure 3.**
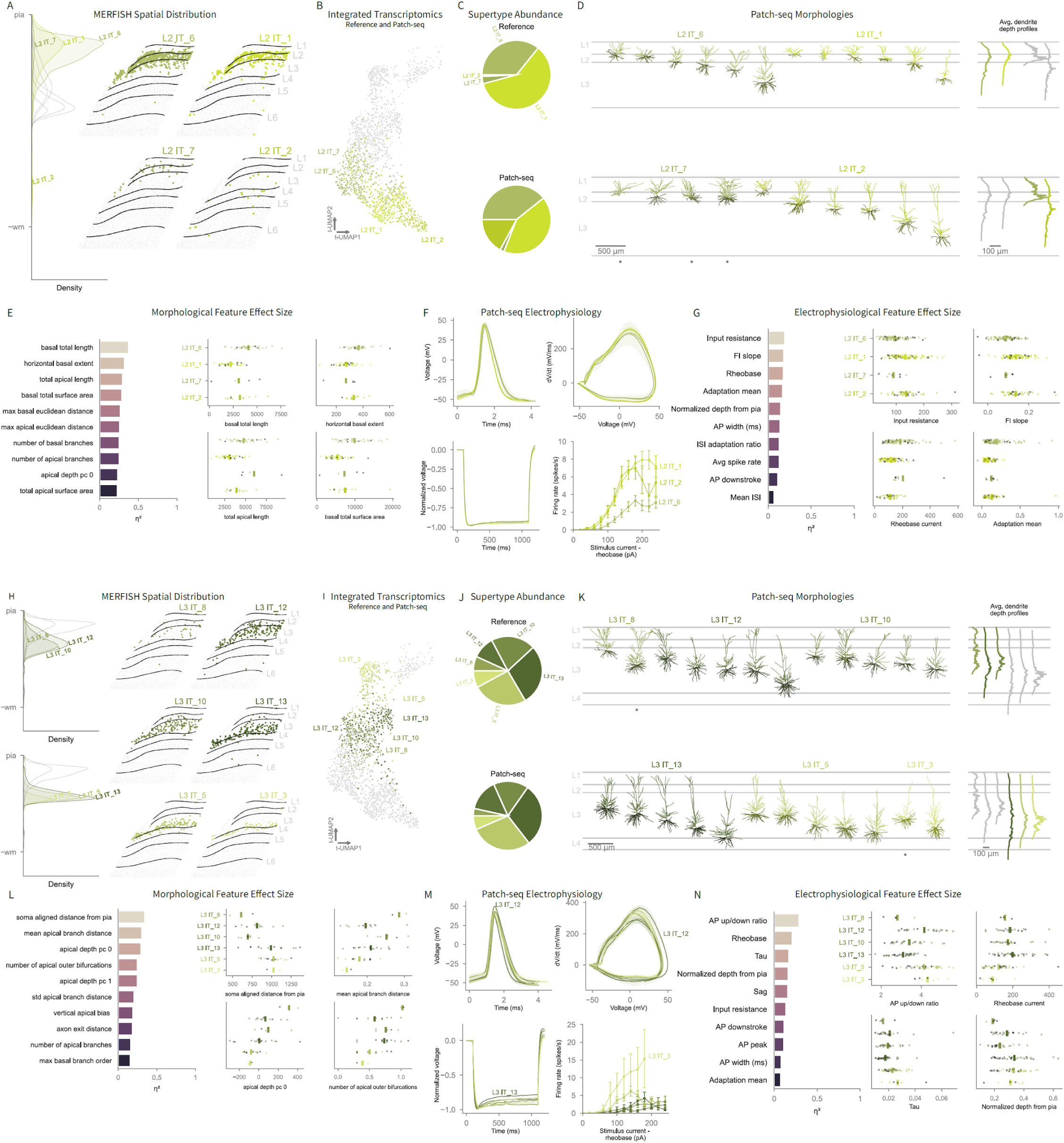
Morphoelectric properties of supertypes in L2 and L3. A. L2 spatial transcriptomics density plot (kernel density estimate) and corresponding spatial transcriptomic sections color coded by supertype. B. Zoomed in view of the L2/3 IT portion of the integrated transcriptomic UMAP in Figure 1D, showing L2 neurons colored by supertype and non L2 IT neurons in grey. C. Proportion of each L2 IT supertype from reference snRNA-seq data (top) and Patch-seq (bottom). D. Representative morphologies from each L2 supertype with histograms (right) showing average dendrite branch length by cortical depth, either colored by supertype or in grey if type is not highlighted in the same row. Grey circle indicates morphologies not from temporal lobe. E. Feature importance analysis of morphological features that best distinguish L2 supertypes. A nonparametric ANOVA (Kruskal-Wallis H test) was used to calculate effect size (η²). Features with η² > 0.4 shown at left and the top four features visualized as strip plots at right colored by supertype; non–temporal lobe samples are shown in grey. F. L2 IT supertype action potentials with corresponding phase plot; voltage response to a 1-s hyperpolarizing current stimulus and the AP firing rate with increasing stimulus intensity above rheobase (f-I) curve averaged by their transcriptomically defined supertype. G. Feature importance analysis of electrophysiological features that best distinguish L2 supertypes. A nonparametric ANOVA (Kruskal-Wallis H test) was used to calculate effect size (η²). Features with η² > 0.4 shown at left and the top four features visualized as strip plots at right colored by supertype; non–temporal lobe samples are shown in grey, culture denoted as open circles and acute as closed circles. H. L3 IT spatial transcriptomics density plot (kernel density estimate) and corresponding spatial transcriptomic sections color coded by supertype. I. Zoomed in view of the L2/3 IT portion of the integrated transcriptomic UMAP in Figure 1D, showing L3 neurons colored by supertype and non L3 IT neurons in grey. J. Proportion of each L3 IT supertype from reference snRNA-seq data (top) and Patch-seq (bottom). K. Representative morphologies from each L3 supertype with histograms (right) showing average dendrite branch length by cortical depth, either colored by supertype or in grey if type is not highlighted in the same row. Grey circle indicates morphologies not from temporal lobe. L. Feature importance analysis of morphological features that best distinguish L3 supertypes. A nonparametric ANOVA (Kruskal-Wallis H test) was used to calculate effect size (η²). Features with η² > 0.4 shown at left and the top four features visualized as strip plots at right. M. L3 IT supertype action potentials with corresponding phase plot; voltage response to a 1-s hyperpolarizing current stimulus and the AP firing rate with increasing stimulus intensity above rheobase (f-I) curve averaged by their transcriptomically defined supertype. N. Feature importance analysis of electrophysiological features that best distinguish L3 supertypes. A nonparametric ANOVA (Kruskal-Wallis H test) was used to calculate effect size (η²). Features with η² > 0.4 shown at left and the top four features visualized as strip plots at right colored by supertype; non–temporal lobe samples are shown in grey, culture denoted as open circles and acute as closed circles.

L2/3 IT_2, which grouped with other L2 supertypes in the integrated transcriptomic UMAP (Figure 3B), had no clear laminar restriction in the MERFISH data (Figure 3A) but was predominately sampled in L2 and L3 with Patch-seq (Figure 1A). Though this supertype appeared to be rare in the MERFISH data, it was overrepresented in Patch-seq relative to the SEA-AD reference dataset (Figure 3C). Using all three modalities to group cells into types^16^, we found that L2/3 IT_2 grouped with L2 IT_1 (Figure S5B). Additionally, many L2/3 IT_2 neurons mapped to a L2 and upper L3 transcriptomics type (GLP2R) identified in our previous study (Figure S6)^1,16,33^. For these reasons, we grouped L2/3 IT_2 with the L2-enriched types. Notably, L2/3 IT_2 is the only L2/3 IT supertype not vulnerable in Alzheimer’s disease^27^.

Morphologically, L2 types varied in complexity. L2 IT_1 neurons appeared to have fewer dendrites than L2 IT_6 (Figure 3D). L2 IT_1 had significantly less apical and basal total length than L2 IT_6 (Figures 3E and S5A). Similarly, L2 IT_1 and L2 IT_2 differed from the other L2 types in their higher input resistance, increased excitability, and steeper F-I slope (Figures 3F, 3G and S5A). When we tested how well a random forest classifier could predict L2 supertypes using either E, M or ME features, the model resulted in 67, 67 and 63% accuracy, respectively, with L2 IT_1 and 6 predicted most reliably (Figure S7). Overall, the L2/3 IT subclass spanned a broad cortical depth range, but several supertypes were enriched in L2, suggesting the need for further refinement of the transcriptomic taxonomy.

### Morphoelectric properties of supertypes in L3

The remaining L2/3 IT supertypes were primarily distributed across L3. Based on the spatial transcriptomic data, supertypes L2/3 IT_8, 10 and 12 were found in upper and mid L3, while L2/3 IT_3, 5, 13 were found in deep L3 (Figure 3H). These supertypes are subsequently referred to as “L3” supertypes.

In the integrated transcriptomic UMAP, the L3 supertypes formed a gradient with superficially enriched supertypes on the south end and deeper supertypes on the north end nearest L4 IT neurons. This spatial arrangement confirmed that cortical depth was strongly encoded in gene expression for all L2/3 supertypes^1,16,19,34,35^ (Figure 3B and 3I).

Morphologically, soma location and apical branching features e.g., mean apical branch distance from soma and the number of outer apical branches, accounted for much of the variation between L3 IT supertypes (Figures 3K and 3L). Electrophysiologically, the action potential kinetics-related feature, up/down ratio, had the largest effect on supertype variance. L3 IT_3, 5 and 12 had a significantly higher up/down stroke ratio, while L3 IT_13 and L3 IT_8 had the lowest (Figures 3M and 3N, Table S2).

A depth-dependent gradient among L2/3 IT neurons has previously been reported^1,34,36^. We revisited these findings using the current, expanded Patch-seq dataset mapped to ten L2/3 IT supertypes, and each assigned to their primary cortical layer. Again, we identified many morphological features, as well as one electrophysiological feature (sag), that significantly varied with depth and/or layer. Furthermore, 21 of 52 morphological features had a significant depth–layer interaction, indicating that the relationship to cortical depth differed between L2 and L3. For example, in L2, the apical dendrite branched further from the soma with increasing depth, whereas the opposite was true for L3. Many morphological features e.g., the number of basal dendritic branches, exhibited steeper depth-related slopes in L2 compared to L3, demonstrating that depth-dependent changes in morphology were more pronounced in L2-dominant supertypes (Figure S5C, Table S2).

### Morphoelectric properties of supertypes in L4

As the major recipient of thalamic and other feedforward inputs, L4 integrates information from many sources^23^. This integrative role suggests that human L4 IT supertypes may also have diverse morphoelectric properties, similar to L4 neurons in mouse somatosensory cortex, where stellate and pyramidal neurons also have distinct afferent connectivity^37^.

Human L4 IT neurons were split into four supertypes that were more discriminable by ME features than any other subclass (Figures 2G and 2H). L4 IT_2 was the major supertype, comprising 75% of the subclass in SEA-AD snRNA-seq reference data, although Patch-seq sampling was closer to 50% (Figure 4A). Laminar position varied slightly across supertypes, with L4 IT_3 and _4 located more superficially than L4 IT_2, while L4 IT_1 was the deepest (Figure 4B). In the integrated transcriptomic UMAP, L4 IT_1 aligned with L5 IT supertypes rather than with other L4 IT supertypes (Figure 4C). Consistent with this, L4 IT_1 and L5 IT_2 supertypes grouped into the same type when all three data modalities were considered, indicating that they shared similar transcriptomics, electrophysiological and morphological properties^16^ (Figure S8A).

**Figure 4.**
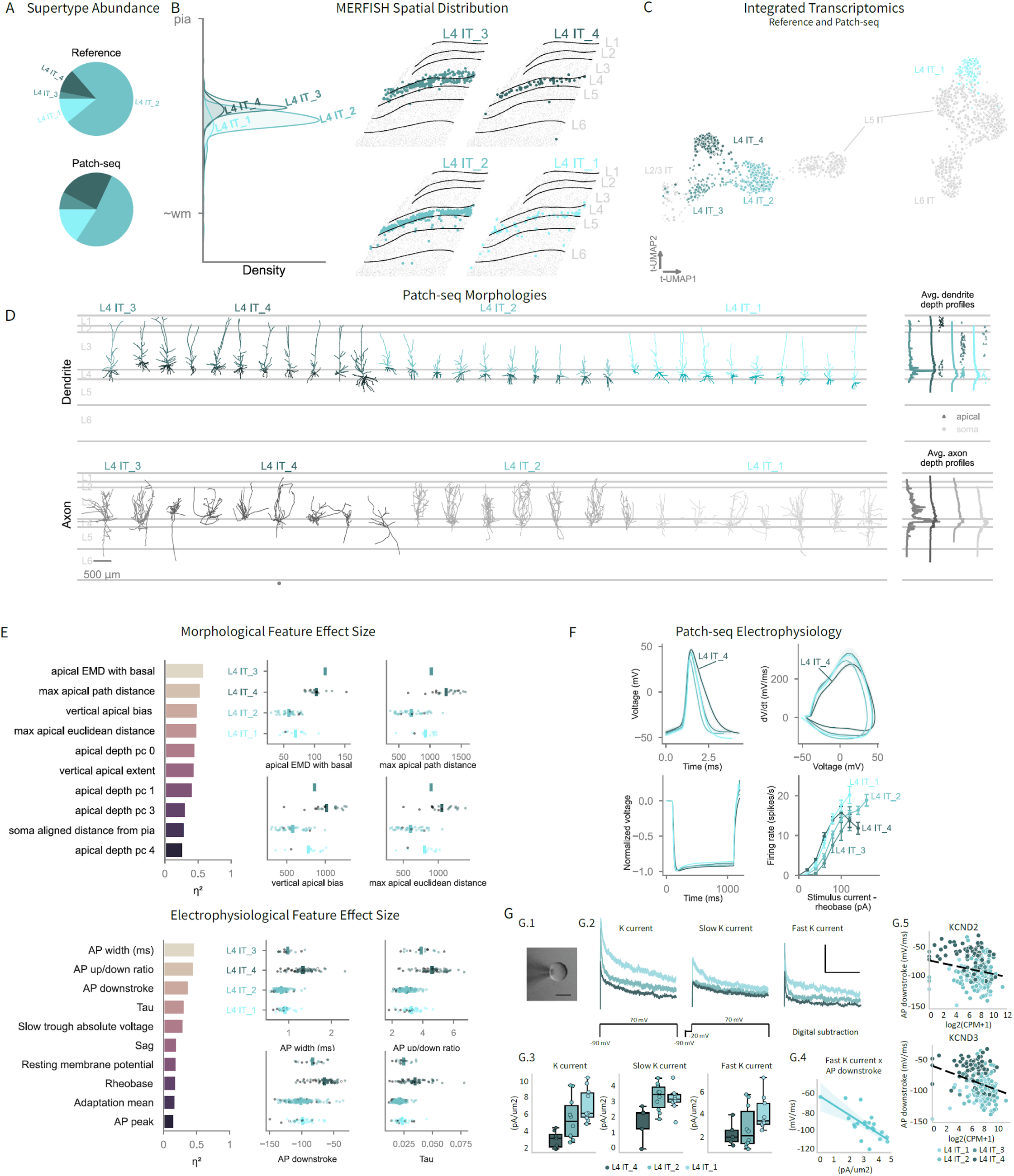
Morphoelectric properties of supertypes in L4. A. Proportion of each L4 IT supertype from reference snRNA-seq data (top) and Patch-seq (bottom). B. L4 IT spatial Transcriptomics density plot (kernel density estimate) and corresponding spatial transcriptomic sections color coded by supertype. C. Zoomed in view of the L4 IT portion of the integrated transcriptomic UMAP in Figure 1D, showing L2 neurons colored by supertype and non-L2 IT neurons in grey. D. Representative morphologies for each supertype. Right: histograms of average neurite length, with apical dendrite tips nearest to layer 1 (triangles) and somas (circles) indicated. Grey circles indicate morphologies not from temporal lobe. E. Feature importance analysis of morphological and electrophysiological features that best distinguish subclasses. A nonparametric ANOVA (Kruskal-Wallis H test) was used to calculate effect size (η²). Features with η² > 0.4 shown at left and the top four features visualized as strip plots at right colored by supertype; non–temporal lobe samples are shown in grey, culture denoted as open circles and acute as closed circles (electrophysiology only). F. L4 IT supertype action potentials with corresponding phase plot; voltage response to a 1-s hyperpolarizing current stimulus and the AP firing rate with increasing stimulus intensity above rheobase (f-I) curve averaged by their transcriptomically defined supertype. G. G.1 Brightfield image of nucleated patch configuration, Scale bar = 5um. G.2 Example total K-current, I_K_ and I_A_ current traces from a +70 mV voltage step for L4 IT_1, L4 IT_2 and L4 IT_2 supertypes, Scale bar = 300 pA/0.5s. G.3 Box and scatter plot depicting the total K-current, I_K_ and I_A_ current for L4 IT_4, L4 IT_2 and L4 IT_1 supertypes. Data is plotted as current density – normalized to size of nucleated patch. G.4 Scatter plot of AP downstroke speed versus I_A_ current for L4 IT subclass nucleated voltage-clamp Patch-seq experiments. G.5 Scatter plot of AP downstroke speed versus KCND2 and KCND3 gene expression for all L4 IT Patch-seq experiments.

The morphologies of the four L4 IT supertypes were quite variable. All four types contained neurons with a prominent apical dendrite with little to no apical tuft (Figure 4D). Morphological features assessed with the d’ metric suggested that L4 IT_4 was the most discriminable type within the L4 IT subclass (Figure S8B). It contained multiple neurons with apical dendrites that reached L1, resulting in greater apical path and Euclidean distance than other L4 IT supertype (Figure 4E). Similar variation has been observed for L4 pyramidal neurons in other cortical regions^8^. Additionally, the local axon of L4 IT_4 differed from L4 IT_2, with L4 IT_2 exhibiting significantly stronger bias towards pia and more bifurcations (Figures 4D and S8C). Based on their high abundance, simple dendrites, and columnar axonal organization, L4 IT_2 neurons were consistent with the canonical thalamocortical-receiving L4 cell type^38,39^.

Electrophysiologically, L4 IT_4 was also the most discernable within the L4 IT subclass (Figures S8B and S8D). Previously described as RORB COL22A1^1^ (Figure S6), L4 IT_4 was distinct from L2/3 IT types and here we show that its electrophysiology properties also differentiated it from other L4 IT supertypes. The average AP waveforms for each supertype revealed that L4 IT_4 exhibited a noticeably broader AP relative to other supertypes (Figure 4F) and the top three distinguishing features were related to AP kinetics (Figure 4E). The variability of the AP width and downstroke among L4 IT supertypes, suggested differential compliments of voltage-gated potassium (K^+^) channels. To assess potential differences in K^+^ channel contributions, we measured perisomatic macroscopic currents in the nucleated patch configuration for a subset of Patch-seq experiments (Figure 4G.1). The L4 IT_4 supertype exhibited the least outward current while L4 IT_1 had the most (Figure 4G.2 and 4G.3). Interestingly, although L4 IT_1 and L4 IT_2 had equivalent sustained current, L4 IT_1 also had the most rapidly inactivating A-type K^+^ current. This current was correlated with the speed of the AP downstroke (Figure 4G.4). To further corroborate this finding, we evaluated gene expression of the canonical A-type K^+^ channel subunits Kv4.2 and Kv4.3 (KCND2 and KCND3) in the larger Patch-seq dataset. Indeed, we observed a significant gene-to-function relationship for both KCND2 and KCND3 with AP downstroke speed (Figure 4G.5), thus providing a functional explanation for the diversity of AP downstroke speeds in L4 IT supertypes.

### Morphoelectric properties of supertypes in L5

L5 is a heterogeneous layer consisting of diverse transcriptomic and axonal projection-defined subclasses. Transcriptomically, glutamatergic subclasses are split into two major groups, IT and non-IT^17^. In L5, the non-IT group contains long-range projecting L5 ET neurons and near projecting L5/6 NP neurons^18,40^. Though L5/6 NP neurons only project within the telencephalon, they are transcriptomically and developmentally related to the non–IT group^21,22^. In total, L5 contains three subclasses, L5 IT, L5 ET and L5/6 NP, and 13 supertypes. In the current analysis, supertypes from the L5 IT subclass corresponded to 85% of all neurons in this layer, while L5/6 NP supertypes accounted for 13.4% and L5 ET for 1.6% (Figure 5A). Within the integrated transcriptomic UMAP, L5/6 NP (5 supertypes) and L5 ET (2 supertypes), formed distinct islands while the six L5 IT supertypes spanned two separate islands with either L4 or L6 IT supertypes as neighbors (Figure 5B).

**Figure 5.**
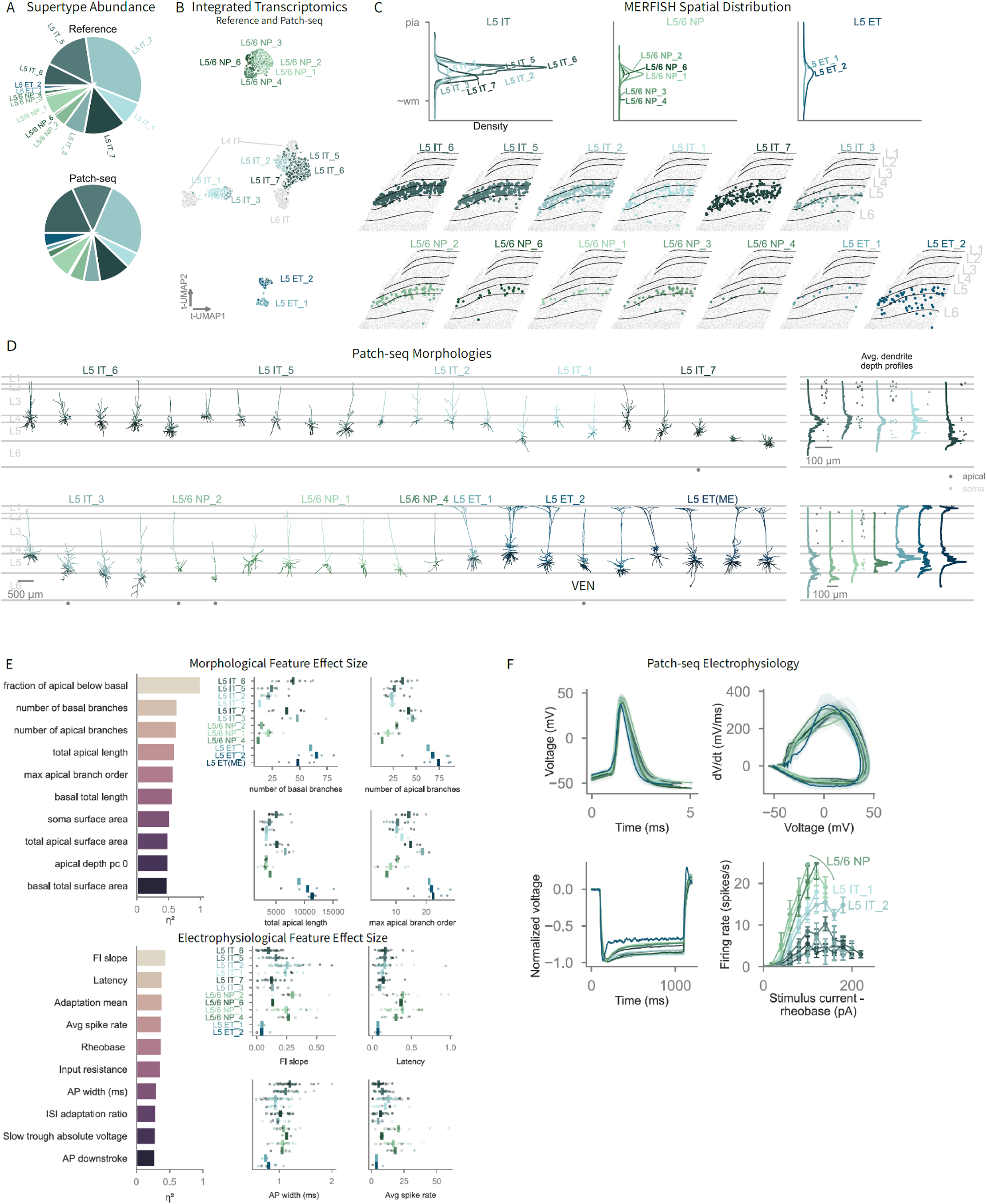
Morphoelectric properties of supertypes in L5. A. Proportion of each L5 supertype from reference snRNA-seq data (top) and Patch-seq (bottom). B. Zoomed in view of the L5 portion of the integrated transcriptomic UMAP in Figure 1D, showing L5 neurons colored by supertype and non L5 neurons in grey. C. L5 spatial transcriptomics density plot (kernel density estimate) and corresponding spatial transcriptomic sections color coded by supertype. D. Representative morphologies for each supertype. Right: histograms of average neurite length, with apical dendrite tips nearest to layer 1 (triangles) and somas (circles) indicated. Grey circles indicate morphologies not from temporal lobe. Von Economo neuron (VEN) is labeled in the plot. E. Feature importance analysis of morphological and electrophysiological features that best distinguish subclasses. A nonparametric ANOVA (Kruskal-Wallis H test) was used to calculate effect size (η²). Features with η² > 0.4 shown at left and the top four features visualized as strip plots at right colored by supertype; non–temporal lobe samples are shown in grey, culture denoted as open circles and acute as closed circles (electrophysiology only). F. L5 supertype action potentials with corresponding phase plot; voltage response to a 1-s hyperpolarizing current stimulus and the AP firing rate with increasing stimulus intensity above rheobase (f-I) curve averaged by their transcriptomically defined supertype.

Given the substantial heterogeneity in L5 (Figures 5A-C and 2H), we explored how well morphological, electrophysiological, and morphoelectric properties can predict subclass and supertype. For subclass, electrophysiological features were highly accurate (96% accuracy; n=243 cells), while morphological features (86%; n=68 cells) and morphoelectric features (92%; n=56 cells) were slightly less so, demonstrating clear subclass boundaries. In contrast, supertype predictions were weaker (54-60% accuracy), suggesting additional heterogeneity at the supertype level (Figure S9A).

The laminar distributions of L5 supertypes overlapped, though supertypes L5 IT_3, _7, L5 ET_2 and L5/6 NP_3, _4 had peak density deeper in L5 (Figure 5C). Most L5 IT types had typical pyramidal characteristics, but L5 IT_3 and 7 had neurons with short apical dendrites, including some stellate-like neurons (Figure 5D). Dendritic branch number and length varied by supertype. NP neurons had relatively sparse dendritic morphologies, while L5 ET neurons had the most extensive branching (Figure 5E). Overall, we found that L5/6 NP neurons had a higher average spike rate and a steeper F-I slope, with a more excitable membrane. Electrophysiologically, NP supertypes were poorly distinguishable from one another (Figure 2G) and using morphology and electrophysiology features in a classifier, they were occasionally predicted as IT supertypes (Figure S9A), highlighting the important role of multimodal features in cell type classification.

L5 ET neurons were separated by a faster AP, higher rheobase, and canonical features such as higher resonance frequency and burst firing^4,41^. Despite their prominent place in the cortical literature, in human cortex, these neurons are rare^4,15^. To supplement the current Patch-seq dataset, we included 23 L5 ET patch-clamp neurons from a previous manuscript^4^ and listed them as “L5 ET(ME)”. Of the two ET supertypes, L5 ET_2 was the most prevalent (71%). Despite transcriptomic differences separating the two L5 ET supertypes, we did not identify any electrophysiological differences, and morphological data for the L5 ET_1 supertype was too limited for a meaningful comparison. L5 ET_2 neurons had diverse apical branching patterns, from main branches that emerged near the soma, to those that branched closer to the apical tuft. A transcriptomically-defined L5 ET neuron sampled from the insular cortex had morphological characteristics of a Von Economo neuron (VEN), including a spindle or fusiform-shaped soma, one large primary basal dendrite and an axon originating from the basal dendrite more than 70 μm from the center of the soma^42–44^. Recent work reported that VENs are regionally specialized ET-projecting neurons^45^; here, we confirmed that a morphologically defined insular VEN mapped to an ET transcriptomic type (Figure S9B).

L5 IT neurons had lower sag ratios compared to L5 ET and L5/6 NP neurons; however, other features within this subclass were more variable. At the supertype level, electrophysiological features varied amongst L5 IT neurons (Figures 2G and 5E). Most notably, L5 IT_1 and L5 IT_2 had steeper F-I slopes like L5/6 NP neurons, while other supertypes had a modest slope (Figure 5F). Morphologically, L5 IT_1 and L5 IT_2 had sparse dendrites with limited apical and basal overlap compared to other L5 IT supertypes, resembling the L4 IT_1 and L4 IT_2 supertypes. L5 IT_2 also had the fastest AP and a higher average spike rate amongst L5 ITs, clustering with L4 IT_1 based on multimodal properties. Additionally, L5 IT_5 and 6 grouped together, while deeper types L5 IT_7 and 3, merged with L6 IT neurons based on their multimodal properties (Figure S9C). These patterns agree with previous findings^4^ that identified two IT-like groups with electrophysiological properties broadly corresponding to SEA-AD supertypes L5 IT_5/7 (IT-like 1) and L5 IT_1/ 2 (IT-like 2). L5 IT_3 neurons, located in deep L5, were not described in that study likely due to its deeper laminar position. Altogether, this analysis revealed complex heterogeneity within L5 IT supertypes, suggesting these prevalent neurons may have more differentiated roles in the human cortical network than previously recognized.

### Morphoelectric properties of supertypes in L6

L6 contains four glutamatergic subclasses, L6 CT, L6 IT, L6 IT Car3 and L6b, more than any other cortical layer. With its cortical and subcortical targets, L6 is thought to have a role in feedback, gain control, and thalamic modulation^46,47^. In this study, we found that L6 had multiple, modified/non-traditional pyramidal neurons, including stellate and bipolar fusiform neurons. While there were 15 supertypes, most were rare, and only a handful accounted for the majority of neurons in L6. In each subclass, a single supertype dominated (L6 CT_2, L6 IT_2, L6b_6) except for L6 IT Car3, where L6 IT Car3_2 and _3 had similar proportions (Figure 6A). L6 supertypes were mainly found within L6, though some extended to L5 or white matter. L6 IT Car3_1 was found closer to and overlapping with L5, which is consistent with this subclass sometimes being referred to as L5/6 Car3 IT^17,19^. In contrast, supertypes like L6 CT_4, _1, and most L6b supertypes, extended into deep L6 and white matter (Figure 6B).

**Figure 6.**
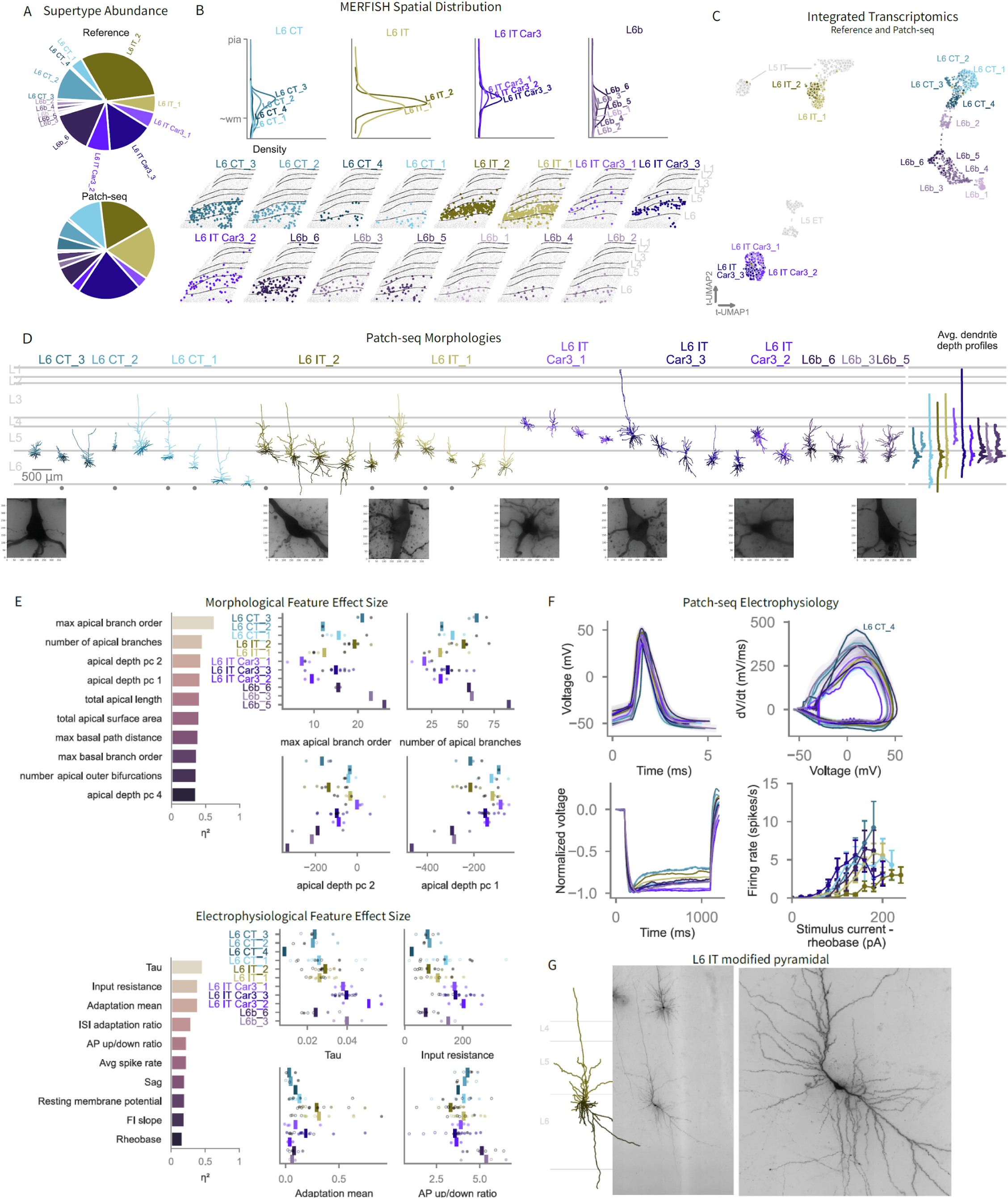
Morphoelectric properties of supertypes in L6. A. Proportion of each L6 supertype from reference snRNA-seq data top and Patch-seq bottom. B. L6 spatial transcriptomics density plot (kernel density estimate) and corresponding spatial transcriptomic sections color coded by supertype. C. Zoomed in view of the L6 portion of the integrated transcriptomic UMAP in Figure 1D, showing L6 neurons colored by supertype and non L5 neurons in grey. D. Representative morphologies from each supertype with histograms (right) showing average dendrite branch length by cortical depth. Grey circle indicates morphologies not from temporal lobe. Perisomatic minimum intensity projections shown below highlighting the unique soma and branching structure of L6 IT supertypes. E. Feature importance analysis of morphological and electrophysiological features that best distinguish subclasses. A nonparametric ANOVA (Kruskal-Wallis H test) was used to calculate effect size (η²). Features with η² > 0.4 shown at left and the top four features visualized as strip plots at right colored by supertype; non–temporal lobe samples are shown in grey, culture denoted as open circles and acute as closed circles (electrophysiology only). F. L6 supertype action potentials with corresponding phase plot; voltage response to a 1-s hyperpolarizing current stimulus and the AP firing rate with increasing stimulus intensity above rheobase (f-I) curve averaged by their transcriptomically defined supertype. G. Reconstruction of a non-traditional/modified L6 IT_2 pyramidal neuron with “tap-root” basal dendrite (left), with high resolution overview (middle) and a zoom in of the perisomatic region.

In the integrated transcriptomic UMAP, L6 IT supertypes were on a separate island near L5 IT supertypes, while L6 IT Car3 supertypes formed a distinct island (Figure 6C). L6 CT and L6b each formed their own islands, and were mostly non-overlapping, with the exception of L6b_2, a rare supertype located in deep L6/white matter that linked the two. Unfortunately, this type was not sampled with Patch-seq.

Most of the non-traditional/modified pyramidal neurons^48^ in this dataset belonged to the L6 IT and L6 IT Car3 subclasses. In both mouse and human, L6 IT Car3^7,49^ neurons are stellate in shape, though in the current dataset, a few human L6 IT Car3_3 neurons deviated from this phenotype, with the most notable exception having a traditional apical dendrite that extended to L1 (Figure 6D). Apical dendrite features, and a few basal dendrite features, explained much of the variance between L6 supertypes (Figure 6E).

For electrophysiology, the membrane time constant (tau) and input resistance emerged as the top two most distinguishing features (Figure 6E). L6 IT Car3 had the highest measured tau and input resistance compared to other L6 subclasses. L6 IT supertypes had the highest average AP firing adaptation, indicating early burst and/or strongly adapting firing patterns (Figure 1A). AP kinetics, measured by the upstroke/downstroke ratio also varied amongst subclasses and supertypes, with L6b neurons having the highest ratio (Figure 6F). The size, extent of the dendrites, variance in AP and AP firing patterns support a unique role for L6 in input/output signaling.

We assessed how well L6 subclasses can be predicted from their morphoelectric properties and achieved the highest accuracy with electrophysiology features (88%) and much lower for morphological features (44%). This drop in accuracy was driven by confusion between L6 CT and L6b neurons and between L6 IT Car3 and L6 IT neurons (Figure S10A). L6 CT neurons provide excitation to the thalamus^47^ and in rodents sublaminar diversity in this population has been observed, with the deeper types preferentially targeting posterior thalamus ^50–52^. Similarly, transcriptomic studies find that specific L6 CT subtypes target specific thalamic nuclei^7,18^. Human L6 CT_1 was the deepest of the CT supertypes (Figure 6B) and had the longest apical dendrites, occasionally reaching layer 3. Although morphological differences among the CT supertypes were not significant, likely reflecting the low sample size (Figure 6D), the deep laminar positions of supertypes L6 CT_1 and_4 may reflect functional differences in humans, as it does in the mouse.

As mentioned, the L6 IT subclass had many distinctive morphologies. A subset of these neurons had elongated somas, and non-tufted apical dendrites that terminated in L3 or L5 and had a prominent, extra-long basal dendrites, sometimes referred to as “tap-root” dendrites^53^. Tap root dendrites were common in L6 IT_2, which were more frequently found in temporal lobe samples, and can exceed 1 mm in length (Figures 6D and 6G). Although previously associated with ET projecting Betz neurons in primary motor cortex^15,54–57^, broader sampling with Patch-seq revealed tap-root dendrites are also found in transcriptomically-defined L5 and L6 IT neurons in mouse^7^ (Figure S10B).

The border from human cortex to white matter can be distinct or gradual, depending on cortical area and location along the gyri^58^, unlike the transition in the mouse that is bound by a thin, distinct L6b. Rodent neurons in L6b have heterogeneous projection targets; some remain within the telencephalon, while others also project to the thalamus^18^. We sampled three of the more superficial L6b supertypes (Figure 6B). These neurons had pia-ascending apical dendrites that didn’t reach beyond L4 and exhibited more extensive apical branching than other L6 subclasses. Among all morphology features, apical branching extent (i.e. max apical branch order) and the number of apical branches, accounted for the greatest variance across all L6 supertypes, with L6b neurons often showing the highest values (Figure 6E). Unlike in the mouse^5,7^, we did not sample human L6b neurons that had dendritic morphologies extending horizontally, presumably as a result of a less compact L6b observed in primates. L6b neurons were of particular interest due to their unique responsiveness to Orexin, a neuropeptide involved in driving wake-like brain states^59^. Orexin receptor type 2 expression was enriched in L6b supertypes except for L6b_2 (Figure S10C).

These analyses revealed that human L6 is a heterogeneous cortical layer, encompassing four neuronal subclasses with distinct morphological and electrophysiological properties. Pronounced dendritic variation, from tap-root dendrites to stellate Car3 neurons, combined with electrophysiological diversity, highlighted the functional complexity of this cortical layer.

### Cross-Species

Humans share many cortical features with macaques and mice; however, their evolutionary divergence occurred approximately 23 and 70 million years ago, respectively. Patch-seq enables cross-species comparisons of morphoelectric properties in transcriptomically homologous cell types. We analyzed previously published and newly generated Patch-seq data from neurons in macaque^54,60^ and human temporal cortex^1,4^, and mouse visual cortex^7^ (Figure 7A). Within the same transcriptomic subclasses, macaque morphology closely matched human, with few significant differences (7%), while nearly all mouse-human morphological comparisons (95%) and most (76%) macaque-mouse comparisons were significant (Figure 7B). Where differences between species existed, neurons tended to be larger in macaque. Specifically, in L5 IT neurons, the number of basal dendrite branches and total basal dendrite length, were significantly greater in macaque; in L2/3 IT neurons, multiple basal and apical features were larger in macaque (Figures 7C and S11A). Overall, human and macaque morphologies were comparable and when differences existed, they varied in a subclass-specific manner.

**Figure 7.**
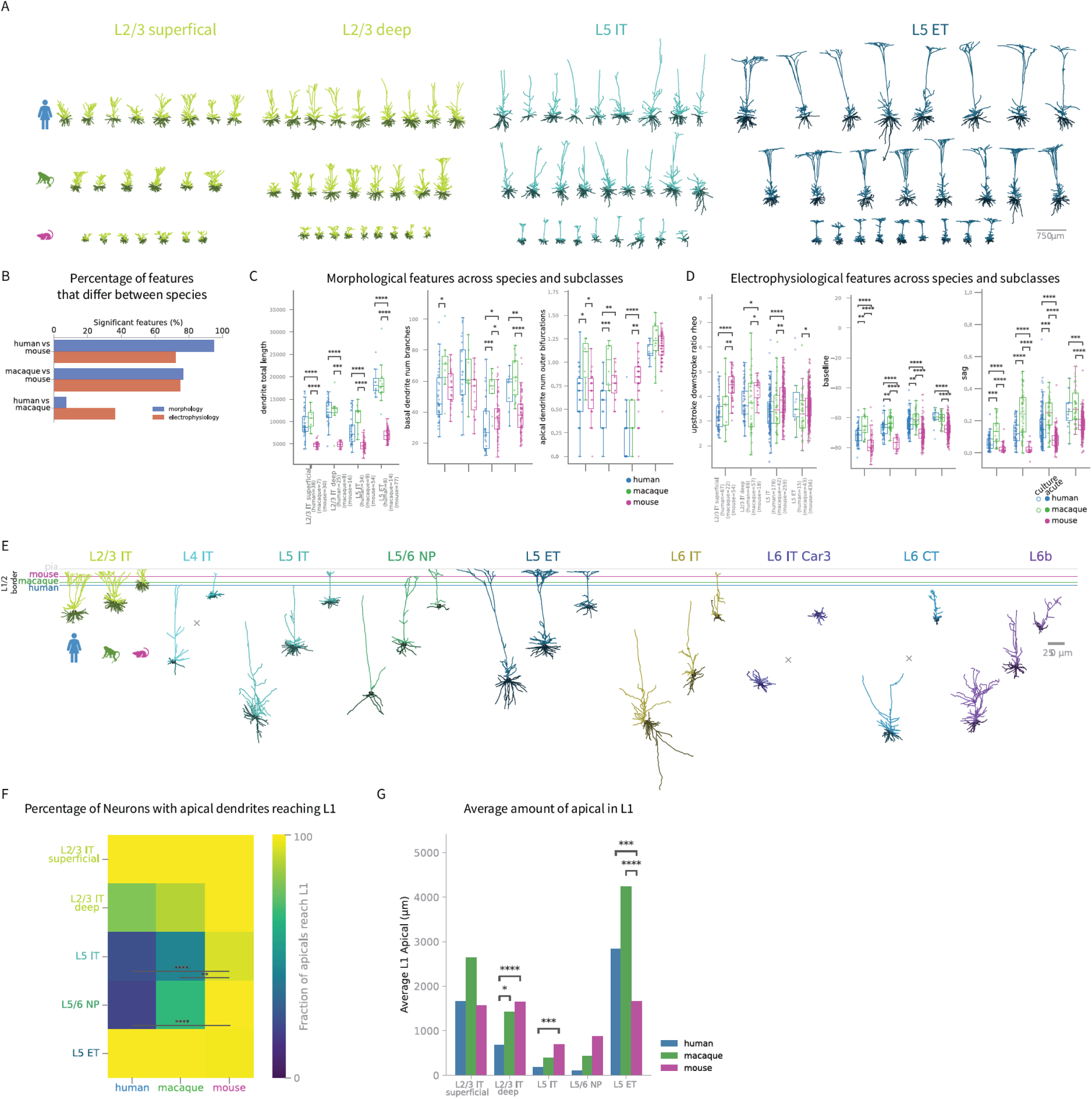
Cross-species analysis of morphoelectric properties of transcriptomic defined subclasses. A. Morphologies aligned by somas across species and subclass. L2/3 IT was broken into two groups, those with somas in the superficial or deep half of layer 2 and layer 3 combined (see methods). B. Percentage of morphological and electrophysiological features that differed between species. C. Morphological feature comparison across species and subclass. Mann-Whitney U-test when only two species groups, otherwise Kruskal-Wallis H-test, FDR corrected. If KW FDR was significant (p < 0.05) post-hoc Dunn test on pairwise species groups, FDR corrected. Full statistical details are provided in Table S2. D. Electrophysiological feature comparison across species and subclass, culture denoted as open circles and acute as closed circles. Mann-Whitney U-test when only two species groups, otherwise Kruskal-Wallis H-test, FDR corrected. If KW FDR is significant (p < 0.05) post-hoc Dunn test on pairwise species groups, FDR corrected. Full statistical details are provided in Table S2. E. Representative morphologies from each subclass across species (human, macaque when available and mouse), aligned to the pia (grey). The L1/2 border from each species’ average layer template is shown in color. F. Heatmap showing the percentage of neurons with apicals reaching L1 across species with significance determined with Chi-square test of independence. Full statistical details are provided in Table S2. G. For neurons with apical dendrites reaching L1, Mann-Whitney U-test (when two species groups) or Kruskal-Wallis H-test (when three species groups), FDR corrected, on the length of apical dendrite in Layer 1 between species groups. If KW FDR is significant (p < 0.05) post-hoc Dunn test on pairwise species groups, FDR corrected. Full statistical details are provided in Table S2.

Next we examined electrophysiological properties across species. Using Linear Discriminate Analysis (LDA) on electrophysiological features for L2/3 IT superficial, L2/3 IT deep, L5 IT and L5 ET subclasses, we identified species-specific differences (Figure S11B). As expected, the most pronounced differences were between primates and mice. In superficial and deep L2/3 IT neurons, humans and macaques exhibited a similar AP up/down ratio that differed from mice. The observed difference in ratio was primarily attributable to a slower AP upstroke in mice. Primates differed in some electrophysical features, where L2/3 (superficial and deep) and L5 IT macaque neurons had a larger sag ratio and more depolarized resting membrane potential compared to their human counterparts (Figure 7D). There were no significant electrophysiology differences between primate L5 ET neurons except the downstroke speed of the AP (Figure S11C). These findings likely reflect distinct, yet complementary contributions of voltage-gated ion channels shaped by subclass identity within each species.

To put cross-species morphoelectric properties in the context of the cortical circuit, we examined apical dendrite patterns with respect to L1. We observed prominent species differences between human and mouse in the amount and frequency of apical dendrites that reach up to L1 (Figure 7E). This agrees with previous work where human infragranular IT neurons were found to be thin-tufted or non-tufted and rarely had apical dendrites that reached L1^4,61,62^, while thick tufted L5 ET neurons had dendrites with strong arborization in L1^4,41^. This differed markedly from the mouse, where both L5 IT and ET neurons frequently had dendritic tufts in L1^5,7,15,30,49^. In the current study, transcriptomically-defined mouse visual area (VIS) L5 IT and L5/6 NP neurons were significantly more likely than human neurons to reach L1 (Figure 7F). Since morphological features vary across cortical regions, increasing in complexity from caudal to rostral^30,63,64^, we also compared human neurons to mouse neurons from rostral brain regions with large total dendrite length ^30,49,65,66^. Projection-defined L5 IT neurons in mouse dorsal and ventral Anterior Cingulate (ACAd/v) and secondary Motor (MOs) cortex were also significantly more likely than human L5 IT neurons to reach L1. This same pattern was also observed for Patch-seq L5 IT neurons in mouse Temporal Association area (TEa) (Figure S11D).

For neurons whose apical dendrites reached L1, we next compared the average amount of apical dendrite in L1 across species and subclasses. In this comparison, mouse neurons had more apical dendrite in L1 than human neurons in deep L2/3 IT and L5 IT subclasses. Deep L2/3 IT macaque neurons also had more apical dendrite in L1 than human but did not significantly differ from mouse. Finally, both human and macaque L5 ET neurons had significantly more L1 apical dendrite than mouse (Figure 7G). These major differences in the propensity of certain subclasses to reach L1 likely translate to important differences in how information is processed in the cortex of different species.

## Discussion

Using Patch-seq, we sampled glutamatergic neurons in human neocortex and mapped them to an existing transcriptomic classification, creating a unified framework for examining the functional properties of 42 glutamatergic transcriptomic supertypes across nine subclasses. We profiled 1491 neurons, primarily in temporal lobe, first analyzing their features at the subclass level and then at the finer granularity of supertype. We systematically characterized this diversity layer-by-layer to provide a unified multimodal reference, supporting experimenters in predicting and identifying cell types in their own studies and enabling modelers to incorporate human-specific morphoelectric and layer-resolved constraints into circuit models. We also provide our dataset in the Cytosplore Viewer, an application for browsing and analyzing the data presented in this manuscript.

Previous studies have illustrated how human supragranular layers have expanded in thickenss^67,68^ and increased transcriptomic cell type diversity^1,16^. In this study, we explored L2/3 IT supertypes relative to their primary layer and found stronger depth-morphoelectric feature correlations in L2 supertypes than in L3, suggesting that proximity to the L1/2 border amplifies depth-dependent effects. These results supported assigning a primary layer to L2/3 IT supertypes. As mentioned, one of the L2/3 IT types (L2 IT_2) was found primarily in L2 and was the only L2 and L3 cell type that was not vulnerable to Alzheimer’s. Differentiating supertypes by their primary layer may facilitate discovery about differential cell type vulnerability in development and disease in these recently expanded layers of human cortex^69–72.^

L4 IT neurons exhibited high input resistance and one of the steepest F-I slopes (second only to L5/6 NP neurons), positioning them as high-gain detectors, particularly well-suited for relaying incoming input. Features of L4 IT_2 neurons, such as their high abundance, simple dendrites, and columnar axon, were consistent with the canonical thalamocortical-receiving L4 cell type^38,39^. L4 IT_4 exhibited a broader AP compared to other L4 IT supertypes and had apical dendrites that regularly reached L1, suggesting it may be important for integrating top-down input via its L1 apical tufts. In addition to L4 serving as a relay for thalamic sensory input, the heterogeneity observed across multimodal features of this population suggests that it may have additional functions.

We presented a novel characterization of human L5/6 NP neurons, a population that may be selectively vulnerable in Huntington’s disease^73^. Similarly, L5 IT and L5 ET neurons are vulnerable in a multitude of disorders^74–76^. Characterization of L5 cell type diversity may help clarify which features underlie their differential vulnerability to disease. We also noted that within certain subclasses, particularly L5/6 NP and L6 IT, supertypes had similar morphoelectric properties. With increased sampling, additional differences may emerge, but it is also possible that their morphoelectric features are orthogonal to their transcriptomic supertype identity.

L6 contained four subclasses with a wide range of morphological and electrophysiological diversity. The axonal signatures of transcriptomically-defined human L6 CT and L6 IT neurons, vertically and horizontally oriented, respectively, matched the contrasting patterns seen in rodent^77^. We also sampled human L6 IT Car3 neurons for the first time, which had mostly compact dendrites with sparse local axon, similar to what has been observed in the mouse^7,49^. Relatedly, membrane properties emerged as the most distinguishing electrophysiological features, reflecting underlying morphological diversity. These findings underscore the likely functional complexity of this layer despite being underexplored in human.

The proportions of L5 IT and L5/6 NP neurons with apical dendrites extending to L1 were different between human and mouse and may suggest that some human glutamatergic neuron subclasses receive less layer 1 input than their mouse counterparts. Top-down integration via L1 input onto the apical tufts of L5 pyramidal neurons is thought to be a key component of cortical circuitry^78^. The observed divergence in L1 apical arborization across species suggests alternative mechanisms for feedback and modulatory input could be at play.

Across the evolutionary spectrum, multimodal features are expected to change in a graded manner as biological systems increase in organization and functional complexity^79^. However, we demonstrated that this pattern is not universal across all subclasses. Macaque morphological and electrophysiological features were often well aligned with human. In certain instances, however, macaque features were distinct, while human and mouse features were more closely aligned. This observation suggests that morphological and electrophysiological features may not adhere to a linear evolutionary pattern but instead appear to vary dynamically across species and to be influenced by transcriptomic identity^60^.

Understanding the heterogeneity of neurons in the human neocortex and how their morphological and electrophysiological features relate to transcriptomic cell type identity is essential for understanding their contributions to circuit function, computational processing^80^, and ultimately cognition. In this study, we provided a comprehensive characterization of human excitatory glutamatergic neurons, which complemented previous work^1–4^, and resulted in the most detailed, multimodal framework for understanding neuronal cell types in the human cortex to date.

Limitations of the Study

Data from all human cortical lobes were grouped in our analysis, which may have obscured region-specific differences. Additionally, some neuron supertypes, especially infragranular types, were under-sampled due to low abundance, highlighting the need for broader and additional sampling across cortical regions. For our cross-species comparisons, the Patch-seq mouse data were collected primarily from sensory cortices (visual), which may not be generalizable to other cortical regions, and are not perfectly matched to many of our human samples, which were primarily obtained from the temporal lobe. To combat this, we also included data from mouse association areas using published single-neuron projectome datasets^30,49,65,66^.

## STAR Methods

### EXPERIMENTAL MODEL AND STUDY PARTICIPANT DETAILS

#### Human neurosurgical specimens and ethical compliance

The neurosurgical tissue specimens collected for this study were apparently non-pathological tissues removed during the normal course of surgery to access underlying pathological tissues. Tissue specimens were determined to be nonessential for diagnostic purposes by medical staff and would have otherwise been discarded. Tissue procurement from neurosurgical donors was performed outside of the supervision of the Allen Institute at a local hospital and tissue was provided to the Allen Institute under the authority of the institutional review board of the participating hospital. A hospital-appointed case coordinator obtained informed consent from the donor before surgery. Tissue specimens were deidentified before receipt by Allen Institute personnel.

#### Non-human primate specimens

We obtained non-human primate brain specimens, Macaca Nemestrina and Macaca Mulatta, from animals designated for the Washington National Primate Research Center’s Tissue Distribution Program. Monkeys were housed in individual cages on a 12h light/dark cycle in a temperature and humidity-controlled room. All procedures involving non-human primates were approved by the University of Washington’s Institutional Care and Use Committee (IACUC) and conformed to the NIH’s Guide for the Care and Use of Laboratory Animals.

## METHOD DETAILS

### Human tissue acquisition

Patch-seq experiments on human neurosurgically resected tissue were conducted at sites in the United States, the Netherlands, and Hungary as part of the BRAIN Initiative Cell Atlas Network (BICAN). Surgical specimens were obtained from local hospitals in collaboration with neurosurgeons. Data included in this study were obtained from neurosurgical tissue resections for the treatment of refractory temporal lobe epilepsy or deep brain tumor. All patients provided informed consent and experimental procedures were approved by hospital institute review boards before commencing the study. Tissue was placed in slicing artificial cerebral spinal fluid (ACSF) as soon as possible following resection. Slicing ACSF comprised (in mM): 92 N-methyl-D-glucamine chloride (NMDG-Cl), 2.5 KCl, 1.2 NaH_2_PO_4_, 30 NaHCO_3_, 20 4-(2-hydroxyethyl)-1-piperazineethanesulfonic acid (HEPES), 25 D-glucose, 2 thiourea, 5 sodium-L-ascorbate, 3 sodium pyruvate, 0.5 CaCl_2_.4H_2_O, and 10 MgSO_4_.7H_2_O. Before use, the solution was equilibrated with 95% O_2_, 5% CO_2_ and the pH was adjusted to 7.3 by addition of 5N HCl solution. Osmolality was verified to be between 295 to 305 mOsm kg^−1^. Human surgical tissue specimens were immediately transported (15 to 35 min) from the hospital site to the laboratory for further processing.

### Tissue processing

Human/macaque acute and cultured brain slices (350 μm) were prepared with a Compresstome VF-300 (Precisionary Instruments) or Leica VT1200S (Leica Biosystems) modified for block-face image acquisition (Mako G125B PoE camera with custom integrated software) before each section to aid in registration to the common reference atlas. Brains or tissue blocks were mounted to preserve intact pyramidal neuron apical dendrites within the brain slice. Slices were transferred to a carbogenated (95% O_2_/5% CO_2_) and warmed (34°C) slicing ACSF to recover for 10 min according to the NMDG protective recovery method^81^. Acute brain slices were then transferred to room temperature holding ACSF of the composition (in mM): 92 NaCl, 2.5 KCl, 1.2 NaH_2_PO_4_, 30 NaHCO_3_, 20 HEPES, 25 D-glucose, 2 thiourea, 5 sodium-L-ascorbate, 3 sodium pyruvate, 2 CaCl_2_.4H_2_O and 2 MgSO_4_.7H_2_O for the remainder of the day until transferred for patch clamp recordings. Before use, the solution was equilibrated with 95% O_2_, 5% CO_2_ and the pH was adjusted to 7.3 using NaOH. Osmolality was verified to be between 295 to 305 mOsm kg^−1^. Alternately, slices for interface culture were placed onto membrane inserts (Millipore) in 6 well plates with 1 mL per well of slice culture media of the composition: 8.4 g/L MEM Eagle medium, 20% heat-inactivated horse serum, 30 mM HEPES, 13 mM d-glucose, 15 mM NaHCO_3_, 1 mM ascorbic acid, 2 mM MgSO_4_·7H_2_O, 1 mM CaCl_2_.4H_2_O, 0.5 mM GlutaMAX-I and 1 mg/L insulin. The slice culture medium was carefully adjusted to pH 7.2 to 7.3 and osmolality of 300 to 310 mOsmoles per kilogram by addition of pure H2O, sterile-filtered and stored at 4°C for up to 2 weeks. Culture plates were placed in a humidified 5% CO_2_ incubator at 35°C, and the slice culture medium was replaced every two to three days until endpoint analysis. One to three hours after brain slices were plated on cell culture inserts, brain slices were infected by direct application of concentrated AAV viral particles over the slice surface^11^.

### AAV vector cloning and viral packaging

Enhancer AAV plasmids were maxi-prepped and transfected with PEI Max 40K (Polysciences Inc., catalog # 24765-1) into one 15-cm plate of AAV-293 cells (Cell Biolabs catalog # AAV-100), along with helper plasmid pHelper (Cell BioLabs) and PHP.eB rep/cap packaging plasmid (Chan *et al*., 2017), with a total mass of 150 μg PEI Max 40K, 30 μg pHelper, 15 μg rep/cap plasmid, and 15 μg enhancer-AAV vector. The next day medium was changed to 1% FBS, and then after 5 days, cells and supernatant were harvested and AAV particles released by three freeze-thaw cycles. Lysate was treated with benzonase after freeze thaw to degrade free DNA (2 μL benzonase, 30 min at 37 degrees, MilliporeSigma catalog # E8263-25KU), and then cell debris was precleared with low-speed spin (1500 g 10 min), and finally the crude virus was concentrated over a 100 kDa molecular weight cutoff Centricon column (MilliporeSigma catalog # Z648043) to a final volume of ∼150 μL. For highly purified large-scale preps this protocol was altered so that ten plates were transfected and harvested together at 3 days after transfection, and then the crude virus was purified by iodixanol gradient centrifugation.

### Patch clamp recording

Slices were continuously perfused (2 mL/min) with fresh, warm (34°C) recording ACSF containing the following (in mM): 126 NaCl, 2.5 KCl, 1.25 NaH_2_PO_4_, 26 NaHCO_3_, 12.5 D-glucose, 2 CaCl_2_.4H_2_O and 2 MgSO_4_.7H_2_O (pH 7.3) and continuously bubbled with 95 % O_2_/5% CO_2_. The bath solution contained blockers of fast glutamatergic (1 mM kynurenic acid) and GABAergic synaptic transmission (0.1 mM picrotoxin). Thick-walled borosilicate glass (Warner Instruments, G150F-3) electrodes were manufactured (Narishige PC-10) with a resistance of 4–5 MΩ. Before recording, the electrodes were filled with ∼1.0 to 1.5 μL of internal solution with biocytin [110 mM potassium gluconate, 10.0 mM HEPES, 0.2 mM ethylene glycol-bis (2-aminoethylether)-N,N,N1,N1tetraacetic acid, 4 mM potassium chloride, 0.3 mM guanosine 51-triphosphate sodium salt hydrate, 10 mM phosphocreatine disodium salt hydrate, 1 mM adenosine 51-triphosphate magnesium salt, 20 μg/mL glycogen, 0.5 U/μL RNAse inhibitor (Takara, 2313A) and 0.5% biocytin (Sigma B4261), pH 7.3]. The pipette was mounted on a Multiclamp 700B amplifier headstage (Molecular Devices) fixed to a micromanipulator (PatchStar, Scientifica). The composition of bath and internal solution as well as preparation methods were chosen to maximize the tissue quality of slices from adult mice, to align with solution compositions typically used in the field (to maximize the chance of comparison to previous studies), modified to reduce RNAse activity and ensure maximal gain of mRNA content.

Electrophysiology signals were recorded using an ITC-18 Data Acquisition Interface (HEKA). Commands were generated, signals processed, and amplifier metadata were acquired using MIES written in Igor Pro (Wavemetrics). Data were filtered (Bessel) at 10 kHz and digitized at 50 kHz. Data were reported uncorrected for the measured^82^ –14 mV liquid junction potential between the electrode and bath solutions.

Prior to data collection, all surfaces, equipment, and materials were thoroughly cleaned in the following manner: a wipe down with DNA away (Thermo Scientific), RNAse Zap (Sigma-Aldrich), and finally nuclease-free water. After formation of a stable seal and break-in, the resting membrane potential of the neuron was recorded (typically within the first minute). A bias current was injected, either manually or automatically using algorithms within the MIES data acquisition package, for the remainder of the experiment to maintain that initial resting membrane potential. Bias currents remained stable for a minimum of 1 s before each stimulus current injection.

To be included in analysis, a neuron needed to have a > 1 GΩ seal recorded before break-in and an initial access resistance <20 MΩ and <15% of the Rinput. To stay below this access resistance cut-off, neurons with a low input resistance were successfully targeted with larger electrodes. For an individual sweep to be included, the following criteria were applied: (i) the bridge balance was <20 MΩ and <15% of the Rinput; (ii) bias (leak) current 0 ± 100 pA; and (iii) root mean square noise measurements in a short window (1.5 ms, to gauge high frequency noise) and longer window (500 ms, to measure patch instability) were <0.07 and 0.5 mV, respectively.

Upon completion of electrophysiological examination, the pipette was centered on the soma or placed near the nucleus (if visible). A small amount of negative pressure was applied (∼–30 mbar) to begin cytosol extraction and attract the nucleus to the tip of the pipette. After approximately one minute, the soma had visibly shrunk and/or the nucleus was near the tip of the pipette. While maintaining the negative pressure, the pipette was slowly retracted in the x and z direction. Slow, continuous movement was maintained while monitoring pipette seal. Once the pipette seal reached >1 GΩ and the nucleus was visible on the tip of the pipette, the speed was increased to remove the pipette from the slice. The pipette containing internal solution, cytosol, and nucleus was removed from the pipette holder and contents were expelled into a PCR tube containing lysis buffer (Takara, 634894).

### SMART-seq v4 RNA-sequencing

The SMART-Seq v4 Ultra Low Input RNA Kit for Sequencing (Takara #634894) was used to reverse transcribe poly(A) RNA and amplify full length cDNA according to the manufacturer’s instructions and as previously described^26^. After reverse transcription, cDNA was amplified with 19 PCR cycles. The NexteraXT DNA Library Preparation (Illumina FC-131-1096) kit with NexteraXT Index Kit V2 Sets A-D (FC-131-2001, 2002, 2003, or 2004) was used for sequencing library preparation. Libraries were sequenced on an Illumina HiSeq 2500 instrument (Illumina HiSeq 2500 System, RRID:SCR_016383) using Illumina High Output V4 chemistry. The following instrumentation software was used during data generation workflow; SoftMax Pro v6.5; VWorks v11.3.0.1195 and v13.1.0.1366; Hamilton Run Time Control v4.4.0.7740; Fragment Analyzer v1.2.0.11; Mantis Control Software v3.9.7.19.

### SMART-seq v4 gene expression quantification

Raw read (fastq) files were aligned to the GRCh38 human genome sequence (Genome Reference Consortium, 2011) with the RefSeq transcriptome version GRCh38.p2 (RefSeq, RRID:SCR_003496, current as of 4/13/2015) and updated by removing duplicate Entrez gene entries from the gtf reference file for STAR processing. For alignment, Illumina sequencing adapters were clipped from the reads using the fastqMCF program (from ea-utils). After clipping, the paired-end reads were mapped using Spliced Transcripts Alignment to a Reference (STAR v2.7.3a, RRID:SCR_015899) using default settings. Reads that did not map to the genome were then aligned to synthetic construct (that is ERCC) sequences and the *E. coli* genome (version ASM584v2). Quantification was performed using summerizeOverlaps from the R package GenomicAlignments v1.18.0. Expression counts were calculated as counts per million (CPM) of exonic plus intronic reads.

### MERFISH data generation

Human postmortem frozen brain tissue was embedded in Optimum Cutting Temperature medium (VWR,25608-930) and sectioned on a Leica cryostat at –17°C at 10 μm onto Vizgen MERSCOPE coverslips. These sections were then processed for MERSCOPE imaging according to the manufacturer’s instructions. Briefly: sections were allowed to adhere to these coverslips at room temperature for 10 min prior to a 1 min wash in nuclease-free phosphate buffered saline (PBS) and fixation for 15 min in 4% paraformaldehyde in PBS. Fixation was followed by three 5 min washes in PBS prior to a 1 min wash in 70% ethanol. Fixed sections were then stored in 70% ethanol at 4C prior to use and for up to one month. Human sections were photobleached using a 150W LED array for 72 hours at 4°C prior to hybridization then washed in 5 ml Sample Prep Wash Buffer (VIZGEN 20300001) in a 5 cm petri dish. Sections were then incubated in 5 ml Formamide Wash Buffer (VIZGEN 20300002) at 37°C for 30 min. Sections were hybridized by placing 50 μl of VIZGEN-supplied Gene Panel Mix onto the section, covering with parafilm and incubating at 37°C for 36 to 48 hours in a humidified hybridization oven. Following hybridization, sections were washed twice in 5 ml Formamide Wash Buffer for 30 min at 47°C. Sections were then embedded in acrylamide by polymerizing VIZGEN Embedding Premix (VIZGEN 20300004) according to the manufacturer’s instructions. Sections were embedded by inverting sections onto 110 μl of Embedding Premix and 10% Ammonium Persulfate (Sigma A3678) and TEMED (BioRad 161-0800) solution applied to a Gel Slick (Lonza 50640) treated 2×3 glass slide. The coverslips were pressed gently onto the acrylamide solution and allowed to polymerize for 1.5h. Following embedding, sections were cleared for 24 to 48 hours with a mixture of VIZGEN Clearing Solution (VIZGEN 20300003) and Proteinase K (New England Biolabs P8107S) according to the Manufacturer’s instructions. Following clearing, sections were washed twice for 5 min in Sample Prep Wash Buffer (PN 20300001). VIZGEN DAPI and PolyT Stain (PN 20300021) was applied to each section for 15 min followed by a 10 min wash in Formamide Wash Buffer. Formamide Wash Buffer was removed and replaced with Sample Prep Wash Buffer during MERSCOPE set up. 100 μl of RNAse Inhibitor (New England BioLabs M0314L) was added to 250 μl of Imaging Buffer Activator (PN 203000015) and this mixture was added through the cartridge activation port to a prethawed and mixed MERSCOPE Imaging cartridge (VIZGEN PN1040004). 15 ml mineral oil (Millipore-Sigma m5904-6X500ML) was added to the activation port and the MERSCOPE fluidics system was primed according to VIZGEN instructions. The flow chamber was assembled with the hybridized and cleared section coverslip according to VIZGEN specifications and the imaging session was initiated after collection of a 10X mosaic DAPI image and selection of the imaging area. For specimens that passed minimum count threshold, imaging was initiated, and processing completed according to VIZGEN proprietary protocol. Following image processing and segmentation, cells with fewer than 50 transcripts are eliminated, as well as cells with volumes falling outside a range of 100 to 300 μm.

The 140-gene Human cortical panel was selected using a combination of manual and algorithmic based strategies requiring a reference single cell/nucleus RNA-seq dataset from the same tissue, in this case the human MTG snRNA-seq dataset and resulting taxonomy^16^. First, an initial set of high-confidence marker genes are selected through a combination of literature search and analysis of the reference data. These genes are used as input for a greedy algorithm (detailed below). Second, the reference RNA-seq data set is filtered to only incude genes compatible with mFISH. Retained genes need to be (i) long enough to allow probe design (>960 base pairs); (ii) expressed highly enough to be detected (FPKM ≥ 10), but not so high as to overcrowd the signal of other genes in a cell (FPKM < 500); (iii) expressed with low expression in off-target cells (FPKM < 50 in nonneuronal cells); and (iv) differentially expressed between cell types (top 500 remaining genes by marker score20). To more evenly sample each cell type, the reference dataset is also filtered to include a maximum of 50 cells per cluster.

The main step of gene selection uses a greedy algorithm to iteratively add genes to the initial set. To do this, each cell in the filtered reference data set is mapped to a cell type by taking the Pearson correlation of its expression counts with each cluster median using the initial gene set of size n, and the cluster corresponding to the maximum value is defined as the “mapped cluster.” The “mapping distance” is then defined as the average cluster distance between the mapped cluster and the originally assigned cluster for each cell. In this case a weighted cluster distance, defined as one minus the Pearson correlation between cluster medians calculated across all filtered genes, is used to penalize cases where cells are mapped to very different types, but an unweighted distance, defined as the fraction of cells that do not map to their assigned cluster, could also be used. This mapping step is repeated for every possible *n* + 1 gene set in the filtered reference data set, and the set with minimum cluster distance is retained as the new gene set.

These steps are repeated using the new get set (of size *n* + 1) until a gene panel of the desired size is attained. Code for reproducing this gene selection strategy is available as part of the mfishtools R library (https://github.com/AllenInstitute/mfishtools).

H5ad creation: Any genes not matched across both the MERSCOPE gene panel and the mapping taxonomy were filtered from the dataset before starting. From there, cluster means were calculated by dividing the number of cells per cluster by the number of clusters collected. Next, we created a training dataset by finding marker genes for each cluster by calculating the l2norm between all clusters and the mean counts of each gene per cluster. This training dataset was fed into a knn alongside the MERSCOPEs cell by gene panel to iteratively calculate best possible gene matches per cluster. All scripts and data used are available at: https://github.com/AllenInstitute/.

### Biocytin histology

During Patch-seq experiments where neurons were filled with biocytin via the patch pipette, brain slices were fixed in 4% paraformaldehyde in 1x phosphate buffered saline (PBS) for 1-2 days followed by PBS at 4⁰ C until staining. A horseradish peroxidase enzyme reaction using diaminobenzidine (DAB) as the chromogen was used to visualize biocytin-filled neurons. Slices were also stained with DAPI and incubated in 1% hydrogen peroxide for 30 minutes. Following permeabilization with 2% Triton-X in PBS, slices were incubated in ABS with 0.1% Triton at 4⁰ C overnight or up to two days and then rinsed slices in PBS three times.

### Imaging of biocytin-labeled neurons

Mounted sections were imaged as described previously.^31^ Slice overview images were captured on either upright AxioImager Z2 microscopes (Zeiss, Germany) equipped with an Axiocam 506 camera and a 20× objective (Zeiss Plan-NEOFLUAR 20x/0.5) as a single-plane DAPI fluorescence and brightfield transmission or on a SLIDEVIEW VS200 (Evident Scientific, Japan) equipped with a Hamamatsu ORCA-Fusion-BT camera and 10× objective (Evident UPLXAPO10X) as a single-plane DAPI fluorescence and brightfield transmission z-stacks (36 – 64 μm in total range, 8 – 12 μm in spacing) with the z-stacks collapsed to minimum intensity projections. After cells were selected from slice overview images, tiled z-stacks of individual cells were acquired in brightfield for the purpose of automated and manual reconstruction using Axioimager Z2 microscopes equipped with 63× objectives (Zeiss LD LCI Plan-Apochromat 63x/1.2 Imm Corr or Zeiss Plan-Apochromat 63x/1.4 Oil) at z-step size of 0.44 μm, or 0.28 μm. Light was transmitted using an oil-immersion objective (1.4 NA). Tiled images were stitched in Zeiss’ ZEN software and exported as individual-plane TIFF files.

### Layer annotation and alignment

To characterize the position of biocytin-labeled cells, a 20× brightfield and fluorescent image of DAPI (41,6-diamidino-2-phenylindole) stained tissue was captured and analyzed to determine layer position. Using the brightfield and DAPI image, soma position and laminar borders were manually drawn for all neurons and were used to calculate depth relative to the pia, white matter, and/or laminar boundaries. Laminar locations were calculated by finding the path connecting pia and white matter that passed through the cell’s soma coordinate, and measuring distance along this path to laminar boundaries, pia and white matter.

For reconstructed neurons, laminar depths were calculated for all segments of the morphology, and these depths were used to create a “layer-aligned” morphology by first rotating the pia-to-WM axis to vertical, then projecting the laminar depth of each segment onto an average cortical layer template.

### Human brain region pinning

Available surgical photodocumentation (MRI or brain model annotation) is used to place the human tissue blocks in approximate 3D space by matching the photodocumentation to a MRI reference brain volume “ICBM 2009b Nonlinear Symmetric”^83^, with Human Common Coordinate Framework overlayed^84^ within the ITK-SNAP interactive software.

### Morphological reconstruction

Reconstructions of apical and basal dendrites and/or axon were generated for neurons with good-quality transcriptomics and biocytin fill. Before selecting a neuron for reconstruction, we examined the apical dendrite (when applicable) in the z-depth of the 63× image stack to confirm it stayed within the slice plane and was not truncated at either slice surface. Axons were similarly prioritized when their trajectory remained largely within the slice plane. Reconstructions were generated based on 63X image stacks described above. Stacks were run through a Vaa3D-based image processing and reconstruction pipeline^85^. An automated reconstruction of some neurons was generated using TReMAP^86^ or the approach described in Gliko 2022^87^. Alternatively, initial reconstructions were created manually using the citizen neuroscience game Mozak (Roskams and Popović, 2016). Automated or manually initiated reconstructions were then extensively manually corrected and curated using a range of tools (for example, virtual finger and polyline) in the Mozak extension (Zoran Popovic, Center for Game Science, University of Washington) of Terafly tools^88,89^ in Vaa3D. After 3D reconstruction, morphological features were calculated (Supplementary Table 3) and also as previously described^6,7,31^.

### Cytosplore Viewer

Exploration of the data is performed by selecting neuron groups of interest, either through associated metadata (e.g., by subclass or supertype) or via free-form brushing within any of the Uniform Manifold Approximation and Projection (UMAP) embeddings. The selected neurons are then simultaneously highlighted across all other visualizations. For those neurons with available morphological reconstructions, their three-dimensional morphologies are rendered at their appropriate cortical depth within the average cortical layer template, alongside a representative electrophysiological voltage trace. This enables straightforward visual comparison between the morphologies of the neurons and their electrophysiological firing patterns. Additional visualizations support numerical feature-importance analysis for both the morphological and electrophysiological features, highlighting which features most strongly distinguish a selected group of neurons from the rest of the dataset. Together, these integrated views enable interactive tri-modal comparison of neuronal groups and facilitate the identification of within- and across-type differences through both visual inspection and quantitative feature analysis. This provides a comprehensive perspective on the characteristics that define specific cell types. This dataset can be viewed and analyzed within the Cytosplore Viewer or downloaded from public data repositories (Figure 1E).

## QUANTIFICATION AND STATISTICAL ANALYSIS

Patch-seq transcriptomic mapping and curation

### Reference transcriptomic dataset

Reference transcriptomic data used in this study were obtained from dissociated nuclei collected from human MTG. We used the “within-species” human MTG reference taxonomy published in Jorstad 2023^17^, and further refined as part of the Seattle Alzheimer’s Disease Brain Cell Atlas (SEA-AD) consortium in Gabitto and Travaglini 2024^27^. These data are publicly accessible at the Allen Brain Map data portal (https://brain-map.org/consortia/sea-ad/human-mtg-10x-sea-ad). In the resulting cell type taxonomy, referred to as the ‘SEA-AD reference taxonomy’ herein, five glutamatergic clusters that existed in the “within-species” human MTG Jorstad 2023^17^ reference were pruned due to high misclassification with other clusters when mapping—L2/3 IT_4, L2/3 IT_9, L2/3 IT_11, L5 IT_4, L5/6 NP_5—leaving a total of 42 Glutamatergic “supertypes”.

### Patch-seq mapping to reference taxonomy

Cells were assigned transcriptomic supertypes using the Hierarchical Nearest Neighbors (HANN) mapping approach^19^, as implemented in the scrattch.mapping R library, as follows. First, we subset the reference taxonomy to exclude non-neuronal types and individually map each Patch-seq cell to this taxonomy, iteratively mapping to class, then subclass, and finally supertype. At each taxonomy level the mapped group is assigned as the minimal correlation distance between the query cell and average expression of cells in each SEA-AD reference taxonomy class/subclass/supertype using pre-calculated sets of marker genes, until the most similar supertype is selected. This iterative process is repeated 100 times per cell using random subsets of marker genes to probabilistically assign supertypes to each neuron: a given neuron’s primary supertype is the supertype mapped to most frequently while alternative transcriptomic supertypes are defined as supertypes mapped to less frequently, if any.

### Reference dataset and Patch-seq transcriptome integration

We defined a quality control metric, using Seurat integration^90^ whereby transcriptomes from the SEA-AD reference taxonomy and the complete set of Patch-seq transcriptomes were aligned and visualized in a Uniform Manifold Approximation and Projection^91^ embedding and performed clustering within this space. We identified a cluster of Patch-seq neurons that did not align with SEA-AD reference taxonomy, many of which also exhibited other metrics of low-quality data (e.g., few genes expressed, high fraction of mitochondrial reads, few UMIs, high glial contamination). To ensure robust analysis of transcriptomic types, these neurons were removed from subsequent analysis.

#### Spatial transcriptomic isodepth calculation

To asses the cortical disruption of supertypes we selected human MTG section, H1930002Cx46MTG202007105, taken from donor, H19.30.002. In this section we selected a polygon region of tissue spanning pia to white matter. This allowed us to sample the full depth of the cortex while avoiding damaged portions of any given section. Cells within this selection with fewer than 10 transcripts were also excluded from further analysis. To estimate the cortical depth of the cells contained within this strip, we used GASTON^92^, a machine learning method that automatically builds a model of cortical depth using gene expression, referred to as the isodepth. We generated this isodepth for the whole section, and visualized the distribution of isodepth for each supertype using kernel density estimation. GASTON parameters can be found in the analysis notebook, available in our repository https://github.com/AllenInstitute/human_patchseq_excitatory.git

#### Electrophysiology feature generation

For all electrophysiology stimuli that elicited spiking, APs were detected by first 40 identifying locations where the smoothed derivative of the membrane potential (dV/dt) exceeded 20 mV ms^−1^, then refining on the basis of several criteria including threshold-to-peak voltage, time differences and absolute peak height. For each AP, threshold, height, width (at half-height), fast after-hyperpolarization (AHP) and interspike trough were calculated (trough and AHP were measured relative to threshold), along with maximal upstroke and downstroke 45 rates dV/dt and the upstroke/downstroke ratio (that is, ratio of the peak upstroke to peak downstroke). Following spike detection, summary features were calculated from sweeps with long square pulse current injection: input resistance (all hyperpolarizing sweeps, –10 to –90 pA), sag (hyperpolarizing sweep with response closest to –100 mV, generally –90 pA stimulus, and depolarizing sag on subthreshold response closest to rheobase), rheobase, and F slope (all five spiking sweeps, up to rheobase +80 pA). Spike train properties were calculated for each spiking sweep: latency, average firing rate, initial instantaneous firing rate (inverse of first ISI), mean and median ISI, ISI CV, irregularity ratio, and adaptation index. These spike train features and the single spike properties listed above (measured on the first AP) were summarized for both the rheobase sweep and a stimulus 40pA above rheobase. For spike upstroke, 10 downstroke, width, threshold, and interspike interval (ISI), “adaptation ratio” features were calculated as a ratio of the spike features between the first and third spike (on the lowest amplitude stimulus to elicit at least four spikes). Spike shape properties were also calculated for short (3 ms) pulse stimulation and a slowly increasing current ramp stimulus (first spike only). A subset of cells also had subthreshold 15 frequency response characterized by a logarithmic chirp stimulus (sine wave with exponentially increasing frequency), for which the impedance profile was calculated and characterized by features including the peak frequency and peak ratio. Feature extraction was implemented using the IPFX python package; custom code used for chirps and some high-level features will be released in a future version of IPFX.

To minimize the influence of subclass-specific feature distributions on imputation, missing electrophysiological values were imputed using a subclass-specific k-nearest neighbors (KNN) approach. For each subclass, a KNN imputer was constructed (using the KNNImputer from the scikit-learn package) using up to 100 neighbors (reduced for small subclasses), and imputation was performed using only cells within the same subclass.

#### Morphology feature generation

Prior to morphological feature analysis, reconstructed neuronal morphologies were expanded in the dimension perpendicular to the cut surface to correct for shrinkage^36,93^ after tissue processing. The amount of shrinkage was calculated by comparing the distance of the soma to the cut surface during recording and after fixation and reconstruction. For primate data, in cases where this shrinkage expansion value was in the 99^th^ percentile of human slices, the average human shrinkage correction value (2.088) was used. Features predominantly determined by differences in the z-dimension were not analyzed to minimize technical artifacts due to z-compression of the slice after processing.

Morphological features were calculated as previously described^31^. In brief, feature definitions were collected from prior studies^94,95^. Features were calculated using the skeleton keys python package (https://github.com/AllenInstitute/skeleton_keys). Features were extracted from neurons aligned in the direction perpendicular to pia and white matter. Laminar axon histograms (bin size of 5 microns) and earth movers distance features require a layer-aligned version of the morphology where node depths are registered to an average laminar depth template.

### Analysis

Unless otherwise specified, statistical analyses of morphological and electrophysiological features were implemented in python using the scipy, scikit_posthocs and statsmodels packages.

### Effect size

To quantify the magnitude of transcriptomic group–related differences in each morphoelectric feature, we computed effect sizes using the eta-squared (η²) statistic derived from the Kruskal–Wallis H test. For each feature, η² reflects the proportion of variance in the ranked data explained by group membership, computed as:

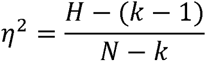

where *H*is the Kruskal–Wallis test statistic, *k* is the number of groups, and *N* is the total sample size. We ordered features by η² to highlight those showing the clearest group-related structure.

Electrophysiology features with values present for fewer than 25% of cells (chirp peak and peak impedance) were excluded from this analysis.

### Random forest classifier

We trained Random Forest classifiers (implemented using the scikit-learn package) to evaluate how well morphology, electrophysiology, and combined morphoelectric features could predict transcriptomic groupings. Transcriptomic groups with at least *n* cells (n=3 within layers; n=4 across layers) were retained. Data were split into an 80/20 stratified train/test set and z-scored using parameters estimated from the training data. Models were fit with class-balanced weights, and hyperparameters were optimized by grid search with 3-fold stratified cross-validation on the training data. The optimal model was refit on the full training set and evaluated on the held-out test set. Accuracy, weighted F1 score, and row-normalized confusion matrices (with the number of test-set cells annotated per row) were computed. Classifiers were trained both with and without cortical depth as a feature, and both unrestricted and restricted to the set of cells with both electrophysiology and morphology data (restricting enables direct comparison across modality classifiers).

### Discriminability (d prime)

Pairwise discriminability (d’) between transcriptomic supertypes was computed within each subclass and cortical layer as previously described^56^. Supertypes with at least three cells were included. Briefly, a Random Forest classifier with balanced class weights was trained on morphoelectric features, and d’ between all supertype pairs were computed from three-fold cross-validated predicted probabilities. Mean d’ values were then calculated by subclass and cortical layer, and displayed with pairwise d’ values for morphology, electrophysiology, and combined morphoelectric feature input.

### Feature box plots

Kruskal–Wallis H tests were performed for each feature across groups of interest (e.g., layers or transcriptomic subclasses). P values were false discovery rate (FDR) corrected using the Benjamini–Hochberg procedure across features, and significant (p<0.05) features were further evaluated with post hoc Dunn tests with FDR correction to identify pairwise group differences.

### Feature x depth regression

To assess how morphological and electrophysiological features vary with cortical depth between layers 2 and 3, we performed a two-way ANOVA including depth from pia, cortical layer, and their interaction as factors, followed by FDR correction using the Benjamini–Hochberg procedure. For selected features, we show the FDR-corrected interaction p-value alongside a scatterplot of layer 2 and layer 3 cells by depth, with within-layer regression lines overlaid (solid if significant, dashed if not).

### MET typing

Electrophysiology features were first reduced to a lower-dimensional representation using sparse principal component analysis (sPCA). The analysis was performed separately on data from each of electrophysiology feature category (e.g., AP waveform, AP features across current steps), and sparse principal components (sPCs) that exceeded 1% adjusted explained variance were kept. This yielded 47 sPCs in total from twelve data categories. The components were z-scored and combined to form the reduced dimension electrophysiology feature matrix used in MET typing.

MET typing was performed as previously described^6^. Briefly, ME clusters and alternative ME-type assignments were generated with random-forest classifiers (90% train, 10% test, bootstrapped 100 times). Combined with a given neurons alternative mapping probability to primary and alternative transcriptomic supertypes we used these cross-mapping probabilities to create a graph of me-type/t-type combinations. A Leiden community detection algorithm^96^ was used to group nodes and generate MET types.

### Apical Reach

To assess subclass differences in apical dendrite reach, we quantified the most superficial apical dendrite position in each reconstructed morphology. We used a Kruskal–Wallis test to determine whether superficial apical dendrite cortical depth differed across subclasses. When the omnibus test was significant (*p* < 0.05), pairwise Dunn’s tests with Benjamini–Hochberg FDR correction were applied to identify subclass-level differences.

### Cross species

#### Cortical depth grouping

To account for differences in cortical depth sampling across species and the merged L2/3 in mouse, L2/3 IT types were subdivided into “superficial” and “deep” groups. Human, macaque, and mouse cells were first aligned by normalized cortical depth, calculated from pia and white-matter fiducials (see *Layer annotation and alignment* methods section). Layer 2 human cells were excluded to match the exclusively layer 3 macaque dataset. Superficial and deep groups were defined using normalized depth thresholds of 0.15–0.25 and 0.25–0.45, respectively, roughly balancing the number of cells from each species.

#### Layer 1 apical reach

To compare apical dendrite presence in layer 1 across species, we first investigated how often apical dendrites made it to layer 1. We extracted the most superficial apical node from neuron reconstructions and measured whether they were within the layer 1 region for that species. For each transcriptomic subclass, we tested whether the proportion of cells with apical dendrites reaching layer 1 differed across species using chi-squared tests. Pairwise Fisher’s exact tests with FDR correction (Benjamini–Hochberg procedure) were applied to identify specific species differences.

For cells with apical dendrite in layer 1, we calculated the length of apical dendrite in layer 1. Subclasses with data from at least two species were tested using a Kruskal–Wallis test (three species) or a Mann–Whitney U test (two species). For subclasses with significant Kruskal–Wallis results (p < 0.05), pairwise Dunn’s tests with Benjamini–Hochberg FDR correction were applied to identify species-level differences. All main-test p-values were subsequently FDR-corrected across subclasses (Benjamini–Hochberg procedure).

#### Species box plots

For each morphoelectric feature, we tested whether values differed across species within each transcriptomic subclass. When data were available for all three species, we applied a Kruskal–Wallis test; if significant (p < 0.05), pairwise differences were assessed using Dunn’s test with Benjamini–Hochberg FDR correction. When only two species had data, we used a Mann–Whitney U test. To control for multiple comparisons across subclasses, all Kruskal–Wallis and Mann–Whitney p-values were subsequently FDR-corrected.

#### Single Neuron Projectome data

CCF-registered whole-neuron morphology reconstructions were aggregated from Gao 2022, Mouselight, Peng 2021, and Castaneda 2021. Dendritic length within cortical layer 1 was quantified by summing the euclidean distances between parent–child node pairs for all dendritic nodes whose coordinates fell within layer 1 of the cortical CCF annotation. To quantify normalized-soma-depth and within-layer-depth we identified the CCF streamline closest to each neuron’s soma (https://github.com/AllenInstitute/ccf_streamlines). For each cell, normalized cortical depth was computed as the distance along this streamline from the soma-nearest point to the pial surface, divided by the total streamline length from pia to white matter. Each voxel in the streamline was annotated with its corresponding cortical layer identity, enabling calculation of depth-within-layer values. Projection patterns were quantified as described in Sorensen et al. 2025^7^. Briefly, a projection matrix was derived based on axonal length within each anatomically defined CCF target structure (https://github.com/matthewMallory/morph_utils). Target regions were evaluated separately in the ipsilateral and contralateral hemispheres. To distinguish true projection targets from fibers of passage, only CCF leaf structures that contained both a branch node and a terminal tip node were included. Long-range projection patterns were used to assign each neuron to a broad projection class (ET, CT, or IT-NP-L6b). Neurons with projections restricted to cortex and cerebral nuclei (CTX/CNU) were labeled IT. Neurons with projections exclusive to cortex were labeled IT-NP-L6b. Neurons with projections extending outside CTX/CNU but limited to thalamus were labeled CT. Neurons with projections to cerebrum and thalamus that also included any projections to hypothalamus, midbrain, hindbrain, or cerebellum were labeled ET.

## Resource availability

### Lead contact

Requests for further information and resources should be directed to and will be fulfilled by the lead contact, Brian R. Lee

### Materials availability

This study did not generate new unique reagents.

### Data and code availability

- *Sequencing data can be found at NEMO:*
- *Electrophysiology data in nwb format have been deposited at DANDI:*
- *Morphological reconstructions in swc format have been deposited at BIL:*
- *All original code has been deposited at GitHub and is publicly available as of the date of publication* https://github.com/AllenInstitute/human_patchseq_excitatory.git

## Acknowledgments

This publication was supported by and coordinated through the BRAIN Initiative Cell Atlas Network (BICAN) (https://braininitiative.nih.gov/research/tools-and-technologies-brain-cells-and-circuits/brain-initiative-cell-atlas-network). This work was funded by the Allen Institute for Brain Science and by the National Institutes of Health awards: UM1 MH130981 (ESL, HZ), R01NS123959 (BEK, ND), U01NS132267 (SAS, TJ) and U01MH114812 (ESL). As well as; The Dutch Research Council (NWO) Open Competition (ENW-M2) grant OCENW. M20.285 (CPdK), Superglue grant number OCENW.XL.23.041 of the research programme ‘NWO Open Competition Domain Science – XL’ from the Dutch Research Council (NWO) (NAG), the European Research Council advanced grant “fasthumanneuron” 101093198 (HDM), Élvonal KKP 133807 (GT) Highlight-152177, Eötvös Loránd Research Network grants ELKH-SZTE Agykérgi Neuronhálózatok Kutatócsoport (GT), KÖ-36/2021 (GT), the NWO Gravitation program BRAINSCAPES: A Roadmap from Neurogenetics to Neurobiology (NWO: 024.004.012) (HDM), TKP-2021-EGA-28 (GT), TKP2021-EGA-09 (GT) and VIDI grant VI.Vidi.213.014 from the Dutch Research Council (NWO) (NAG).

## Author Contributions

Tissue acquisition/processing: AAG, AGA, AJ, AJK, ALK, AP, AT, BG, BRL, CC, CE, DC, DCJ, DLS, DN, EB, EJM, EP, FW, HT, JGO, JGl, JJR, JS, JSH, KC, KW, LE, LW, MC, MF, MJ, MKa, MKr, MRM, NAG, ND, NS, NV, REl, RPG, SE, SI, SLD, SS, TCas, TPS, TSH, TVC, VDM, WH

Transcriptomic processing : CRi, DAM, DB, JAM, KAS, MT, TCar

Spatial transcriptomic data generation: BLo, DAM, EG, JC, JM

Reconstruction: AK, AMu, ÁgKK, CRa, DS, EJM, ÉvT, FW, GW, IR, JAn, LA, RD, RdF, SK

Electrophsiology: AAG, AB, AJK, AMc, AOl, BRL, CRa, DCJ, DH, EJM, FW, GL, JTr, KBa, KBl, KH, KN, LKi, LN, LW, MCV, MKi, MKo, MRM, MV, NAG, NCD, NS, RM, RR, SLD, SS, SV, TPS, TSH, TVC, VDM

Imaging/Histology: AA, AAG, AG, AH, AJK, AOy, AR, ARS, CAP, DCJ, DP, EJM, FW, HG, JAr, JBo, JM, JW, KBe, KBi, KBr, LP, LW, MB, MMax, MMc, MRM, NID, NS, REn, SB, SDH, SLD, STR, TE, TPS, TSH, TVC, VDM, ZCJ,

Data archive / Infrastructure: ABC, BLel, BRL, DR, JGo, JTh, KAS, MMal, MT, NWG, RD, SW

Managment/Project Managment: ABC, BK, BLel, BLo, BRL, CPdK, ESL, HDM, HZ, JW, KAS, LKr, LP, NAG, NCD, NWG, RD, SAS, SDH, TJ

Data analysis: AK, AMc, BRL, EG, FW, JAM, MMal, RD, STR, SW

Data interpretation: BK, BRL, CPdK, ESL, FW, GT, HDM, HZ, JAM, JTT, JTh, NAG, NCD, NWG, RD, SAS, SW, TJ, XL

Writing manuscript: AB, BK, BLel, BRL, CPdK, EG, FW, HDM, JAM, JTT, JTh, MMal, NAG, NCD, RD, SAS, SDH, SW, TJ

Funding acquisition: BK, CPdK, ESL, GT, HDM, HZ, NAG, NCD, SAS, TJ

## Declaration of Generative AI and AI-Assisted Technologies

During the preparation of this work, the authors used OpenAI ChatGPT to debug code and ChatGPT / Microsoft Copilot to improve the readability and language of the manuscript. After using this tool, the authors reviewed and edited the content as needed and take full responsibility for the content of the publication.

## References

1. Berg, J., Sorensen, S.A., Ting, J.T., Miller, J.A., Chartrand, T., Buchin, A., Bakken, T.E., Budzillo, A., Dee, N., Ding, S.-L., et al. (2021). Human neocortical expansion involves glutamatergic neuron diversification. Nature 598, 151–158. 10.1038/s41586-021-03813-8.

2. Lee, B.R., Dalley, R., Miller, J.A., Chartrand, T., Close, J., Mann, R., Mukora, A., Ng, L., Alfiler, L., Baker, K., et al. (2023). Signature morphoelectric properties of diverse GABAergic interneurons in the human neocortex. Science 382, eadf6484. 10.1126/science.adf6484.

3. Chartrand, T., Dalley, R., Close, J., Goriounova, N.A., Lee, B.R., Mann, R., Miller, J.A., Molnar, G., Mukora, A., Alfiler, L., et al. (2023). Morphoelectric and transcriptomic divergence of the layer 1 interneuron repertoire in human versus mouse neocortex. Science 382, 1–18. DOI:%252010.1126/science.adj0805.

4. Kalmbach, B.E., Hodge, R.D., Jorstad, N.L., Owen, S., de Frates, R., Yanny, A.M., Dalley, R., Mallory, M., Graybuck, L.T., Radaelli, C., et al. (2021). Signature morpho-electric, transcriptomic, and dendritic properties of human layer 5 neocortical pyramidal neurons. Neuron 109, 2914–2927.e5. 10.1016/j.neuron.2021.08.030.

5. Scala, F., Kobak, D., Bernabucci, M., Bernaerts, Y., Cadwell, C.R., Castro, J.R., Hartmanis, L., Jiang, X., Laturnus, S., Miranda, E., et al. (2021). Phenotypic variation of transcriptomic cell types in mouse motor cortex. Nature 598, 144–150. 10.1038/s41586-020-2907-3.

6. Gouwens, N.W., Sorensen, S.A., Baftizadeh, F., Budzillo, A., Lee, B.R., Jarsky, T., Alfiler, L., Baker, K., Barkan, E., Berry, K., et al. (2020). Integrated Morphoelectric and Transcriptomic Classification of Cortical GABAergic Cells. Cell 183, 935–953.e19. 10.1016/j.cell.2020.09.057.

7. Sorensen, S.A., Gouwens, N.W., Wang, Y., Mallory, M., Budzillo, A., Dalley, R., Lee, B., Gliko, O., Kuo, H.-C., Kuang, X., et al. (2023). Connecting single-cell transcriptomes to projectomes in mouse visual cortex. bioRxiv, 2023.11.25.568393. 10.1101/2023.11.25.568393.

8. Scala, F., Kobak, D., Shan, S., Bernaerts, Y., Laturnus, S., Cadwell, C.R., Hartmanis, L., Froudarakis, E., Castro, J.R., Tan, Z.H., et al. (2019). Layer 4 of mouse neocortex differs in cell types and circuit organization between sensory areas. Nat Commun 10, 4174. 10.1038/s41467-019-12058-z.

9. Mich, J.K., Graybuck, L.T., Hess, E.E., Mahoney, J.T., Kojima, Y., Ding, Y., Somasundaram, S., Miller, J.A., Kalmbach, B.E., Radaelli, C., et al. (2021). Functional enhancer elements drive subclass-selective expression from mouse to primate neocortex. Cell Reports 34, 108754. 10.1016/j.celrep.2021.108754.

10. Johansen, N.J., Kempynck, N., Zemke, N.R., Somasundaram, S., De Winter, S., Hooper, M., Dwivedi, D., Lohia, R., Wehbe, F., Li, B., et al. (2025). Evaluating methods for the prediction of cell-type-specific enhancers in the mammalian cortex. Cell Genom 5, 100879. 10.1016/j.xgen.2025.100879.

11. Ting, J.T., Kalmbach, B., Chong, P., de Frates, R., Keene, C.D., Gwinn, R.P., Cobbs, C., Ko, A.L., Ojemann, J.G., Ellenbogen, R.G., et al. (2018). A robust ex vivo experimental platform for molecular-genetic dissection of adult human neocortical cell types and circuits. Sci Rep 8, 8407. 10.1038/s41598-018-26803-9.

12. Luo, L., Callaway, E.M., and Svoboda, K. (2008). Genetic Dissection of Neural Circuits. Neuron 57, 634–660. 10.1016/j.neuron.2008.01.002.

13. Zeng, H. (2022). What is a cell type and how to define it? Cell 185, 2739–2755. 10.1016/j.cell.2022.06.031.

14. Jorstad, N.L., Close, J., Johansen, N., Yanny, A.M., Barkan, E.R., Travaglini, K.J., Bertagnolli, D., Campos, J., Casper, T., Crichton, K., et al. (2023). Transcriptomic cytoarchitecture reveals principles of human neocortex organization. Science 382, eadf6812. 10.1126/science.adf6812.

15. Bakken, T.E., Jorstad, N.L., Hu, Q., Lake, B.B., Tian, W., Kalmbach, B.E., Crow, M., Hodge, R.D., Krienen, F.M., Sorensen, S.A., et al. (2021). Comparative cellular analysis of motor cortex in human, marmoset and mouse. Nature 598, 111–119. 10.1038/s41586-021-03465-8.

16. Hodge, R.D., Bakken, T.E., Miller, J.A., Smith, K.A., Barkan, E.R., Graybuck, L.T., Close, J.L., Long, B., Johansen, N., Penn, O., et al. (2019). Conserved cell types with divergent features in human versus mouse cortex. Nature 573, 61–68. 10.1038/s41586-019-1506-7.

17. Jorstad, N.L., Song, J.H.T., Exposito-Alonso, D., Suresh, H., Castro-Pacheco, N., Krienen, F.M., Yanny, A.M., Close, J., Gelfand, E., Long, B., et al. (2023). Comparative transcriptomics reveals human-specific cortical features. Science 382, eade9516. 10.1126/science.ade9516.

18. Tasic, B., Yao, Z., Graybuck, L.T., Smith, K.A., Nguyen, T.N., Bertagnolli, D., Goldy, J., Garren, E., Economo, M.N., Viswanathan, S., et al. (2018). Shared and distinct transcriptomic cell types across neocortical areas. Nature 563, 72. 10.1038/s41586-018-0654-5.

19. Yao, Z., Velthoven, C.T.J. van, Nguyen, T.N., Goldy, J., Sedeno-Cortes, A.E., Baftizadeh, F., Bertagnolli, D., Casper, T., Chiang, M., Crichton, K., et al. (2021). A taxonomy of transcriptomic cell types across the isocortex and hippocampal formation. Cell 184, 3222–3241.e26. 10.1016/j.cell.2021.04.021.

20. Siletti, K., Hodge, R., Mossi Albiach, A., Lee, K.W., Ding, S.-L., Hu, L., Lönnerberg, P., Bakken, T., Casper, T., Clark, M., et al. (2023). Transcriptomic diversity of cell types across the adult human brain. Science 382, eadd7046. 10.1126/science.add7046.

21. Gao, Y., van Velthoven, C.T.J., Lee, C., Thomas, E.D., Mathieu, R., Ayala, A.P., Barta, S., Bertagnolli, D., Campos, J., Cardenas, T., et al. (2025). Continuous cell-type diversification in mouse visual cortex development. Nature 647, 127–142. 10.1038/s41586-025-09644-1.

22. Wang, L., Wang, C., Moriano, J.A., Chen, S., Zuo, G., Cebrián-Silla, A., Zhang, S., Mukhtar, T., Wang, S., Song, M., et al. (2025). Molecular and cellular dynamics of the developing human neocortex. Nature 647, 169–178. 10.1038/s41586-024-08351-7.

23. Harris, K.D., and Shepherd, G.M.G. (2015). The neocortical circuit: themes and variations. Nature Neuroscience 18, 170–181.

24. Zhou, J., Zhang, Z., Wu, M., Liu, H., Pang, Y., Bartlett, A., Peng, Z., Ding, W., Rivkin, A., Lagos, W.N., et al. (2023). Brain-wide correspondence of neuronal epigenomics and distant projections. Nature 624, 355–365. 10.1038/s41586-023-06823-w.

25. Cadwell, C.R., Palasantza, A., Jiang, X., Berens, P., Deng, Q., Yilmaz, M., Reimer, J., Shen, S., Bethge, M., Tolias, K.F., et al. (2016). Electrophysiological, transcriptomic and morphologic profiling of single neurons using Patch-seq. Nat Biotechnol 34, 199–203. 10.1038/nbt.3445.

26. Lee, B.R., Budzillo, A., Hadley, K., Miller, J.A., Jarsky, T., Baker, K., Hill, D., Kim, L., Mann, R., Ng, L., et al. (2021). Scaled, high fidelity electrophysiological, morphological, and transcriptomic cell characterization. eLife 10, e65482. 10.7554/eLife.65482.

27. Gabitto, M.I., Travaglini, K.J., Rachleff, V.M., Kaplan, E.S., Long, B., Ariza, J., Ding, Y., Mahoney, J.T., Dee, N., Goldy, J., et al. (2024). Integrated multimodal cell atlas of Alzheimer’s disease. Nat Neurosci 27, 2366–2383. 10.1038/s41593-024-01774-5.

28. van Unen, V., Höllt, T., Pezzotti, N., Li, N., Reinders, M.J.T., Eisemann, E., Koning, F., Vilanova, A., and Lelieveldt, B.P.F. (2017). Visual analysis of mass cytometry data by hierarchical stochastic neighbour embedding reveals rare cell types. Nat Commun 8, 1740. 10.1038/s41467-017-01689-9.

29. Höllt, T., Pezzotti, N., van Unen, V., Koning, F., Eisemann, E., Lelieveldt, B., and Vilanova, A. (2016). Cytosplore: Interactive Immune Cell Phenotyping for Large Single-Cell Datasets. Computer Graphics Forum 35, 171–180. 10.1111/cgf.12893.

30. Gao, L., Liu, S., Wang, Y., Wu, Q., Gou, L., and Yan, J. (2023). Single-neuron analysis of dendrites and axons reveals the network organization in mouse prefrontal cortex. Nat Neurosci 26, 1111–1126. 10.1038/s41593-023-01339-y.

31. Gouwens, N.W., Sorensen, S.A., Berg, J., Lee, C., Jarsky, T., Ting, J., Sunkin, S.M., Feng, D., Anastassiou, C.A., Barkan, E., et al. (2019). Classification of electrophysiological and morphological neuron types in the mouse visual cortex. Nature Neuroscience, 1. 10.1038/s41593-019-0417-0.

32. Schneider-Mizell, C.M., Bodor, A.L., Brittain, D., Buchanan, J., Bumbarger, D.J., Elabbady, L., Gamlin, C., Kapner, D., Kinn, S., Mahalingam, G., et al. (2025). Inhibitory specificity from a connectomic census of mouse visual cortex. Nature 640, 448–458. 10.1038/s41586-024-07780-8.

33. Miller, J.A., Travaglini, K.J., Luquez, T., Hostetler, R.E., Oster, A., Daniel, S., Tasic, B., and Menon, V. (2025). Annotation Comparison Explorer (ACE): connecting brain cell types across studies of health and Alzheimer’s Disease. Preprint at bioRxiv, 10.1101/2025.02.11.637559 https://doi.org/10.1101/2025.02.11.637559.

34. Kalmbach, B.E., Buchin, A., Long, B., Close, J., Nandi, A., Miller, J.A., Bakken, T.E., Hodge, R.D., Chong, P., Frates, R. de, et al. (2018). h-Channels Contribute to Divergent Intrinsic Membrane Properties of Supragranular Pyramidal Neurons in Human versus Mouse Cerebral Cortex. Neuron 100, 1194–1208.e5. 10.1016/j.neuron.2018.10.012.

35. Cheng, S., Butrus, S., Tan, L., Xu, R., Sagireddy, S., Trachtenberg, J.T., Shekhar, K., and Zipursky, S.L. (2022). Vision-dependent specification of cell types and function in the developing cortex. Cell 185, 311–327.e24. 10.1016/j.cell.2021.12.022.

36. Deitcher, Y., Eyal, G., Kanari, L., Verhoog, M.B., Atenekeng Kahou, G.A., Mansvelder, H.D., de Kock, C.P.J., and Segev, I. (2017). Comprehensive Morpho-Electrotonic Analysis Shows 2 Distinct Classes of L2 and L3 Pyramidal Neurons in Human Temporal Cortex. Cereb Cortex 27, 5398–5414. 10.1093/cercor/bhx226.

37. Schubert, D., Kötter, R., Zilles, K., Luhmann, H.J., and Staiger, J.F. (2003). Cell Type-Specific Circuits of Cortical Layer IV Spiny Neurons. J. Neurosci. 23, 2961–2970. 10.1523/JNEUROSCI.23-07-02961.2003.

38. Staiger, J.F. (2004). Functional Diversity of Layer IV Spiny Neurons in Rat Somatosensory Cortex: Quantitative Morphology of Electrophysiologically Characterized and Biocytin Labeled Cells. Cerebral Cortex 14, 690–701. 10.1093/cercor/bhh029.

39. Narayanan, R.T., Egger, R., Johnson, A.S., Mansvelder, H.D., Sakmann, B., de Kock, C.P.J., and Oberlaender, M. (2015). Beyond Columnar Organization: Cell Type- and Target Layer-Specific Principles of Horizontal Axon Projection Patterns in Rat Vibrissal Cortex. Cereb Cortex 25, 4450–4468. 10.1093/cercor/bhv053.

40. Sorensen, S.A., Bernard, A., Menon, V., Royall, J.J., Glattfelder, K.J., Desta, T., Hirokawa, K., Mortrud, M., Miller, J.A., Zeng, H., et al. (2015). Correlated Gene Expression and Target Specificity Demonstrate Excitatory Projection Neuron Diversity. Cereb Cortex 25, 433–449. 10.1093/cercor/bht243.

41. Beaulieu-Laroche, L., Toloza, E.H.S., van der Goes, M.-S., Lafourcade, M., Barnagian, D., Williams, Z.M., Eskandar, E.N., Frosch, M.P., Cash, S.S., and Harnett, M.T. (2018). Enhanced Dendritic Compartmentalization in Human Cortical Neurons. Cell 175, 643–651.e14. 10.1016/j.cell.2018.08.045.

42. Banovac, I., Sedmak, D., Džaja, D., Jalšovec, D., Jovanov Milošević, N., Rašin, M.R., and Petanjek, Z. (2019). Somato-dendritic morphology and axon origin site specify von Economo neurons as a subclass of modified pyramidal neurons in the human anterior cingulate cortex. J Anat 235, 651–669. 10.1111/joa.13068.

43. Banovac, I., Sedmak, D., Judaš, M., and Petanjek, Z. (2021). Von Economo Neurons – Primate-Specific or Commonplace in the Mammalian Brain? Frontiers in Neural Circuits 15.

44. González-Acosta, C.A., Ortiz-Muñoz, D., Becerra-Hernández, L.V., Casanova, M.F., and Buriticá, E. (2022). Von Economo neurons: Cellular specialization of human limbic cortices? Journal of Anatomy 241, 20–32. 10.1111/joa.13642.

45. Hodge, R.D., Miller, J.A., Novotny, M., Kalmbach, B.E., Ting, J.T., Bakken, T.E., Aevermann, B.D., Barkan, E.R., Berkowitz-Cerasano, M.L., Cobbs, C., et al. (2020). Transcriptomic evidence that von Economo neurons are regionally specialized extratelencephalic-projecting excitatory neurons. Nat Commun 11, 1172. 10.1038/s41467-020-14952-3.

46. Olsen, S.R., Bortone, D.S., Adesnik, H., and Scanziani, M. (2012). Gain control by layer six in cortical circuits of vision. Nature 483, 47–52. 10.1038/nature10835.

47. Crandall, S.R., Cruikshank, S.J., and Connors, B.W. (2015). A Corticothalamic Switch: Controlling the Thalamus with Dynamic Synapses. Neuron 86, 768–782. 10.1016/j.neuron.2015.03.040.

48. Rasia-Filho, A.A., Guerra, K.T.K., Vásquez, C.E., Dall’Oglio, A., Reberger, R., Jung, C.R., and Calcagnotto, M.E. (2021). The Subcortical-Allocortical-Neocortical continuum for the Emergence and Morphological Heterogeneity of Pyramidal Neurons in the Human Brain. Front. Synaptic Neurosci. 13. 10.3389/fnsyn.2021.616607.

49. Peng, H., Xie, P., Liu, L., Kuang, X., Wang, Y., Qu, L., Gong, H., Jiang, S., Li, A., Ruan, Z., et al. (2021). Morphological diversity of single neurons in molecularly defined cell types. Nature 598, 174–181. 10.1038/s41586-021-03941-1.

50. Zhang, Z.-W., and Deschênes, M. (1997). Intracortical Axonal Projections of Lamina VI Cells of the Primary Somatosensory Cortex in the Rat: A Single-Cell Labeling Study. J. Neurosci. 17, 6365–6379. 10.1523/JNEUROSCI.17-16-06365.1997.

51. Bourassa, J., and Deschênes, M. (1995). Corticothalamic projections from the primary visual cortex in rats: a single fiber study using biocytin as an anterograde tracer. Neuroscience 66, 253–263. 10.1016/0306-4522(95)00009-8.

52. Breuer, T.M., and Krieger, P. (2022). Sensory deprivation leads to subpopulation-specific changes in layer 6 corticothalamic cells. European Journal of Neuroscience 55, 566–588. 10.1111/ejn.15572.

53. Scheibel, M.E., Tomiyasu, U., and Scheibel, A.B. (1977). The aging human Betz cell. Exp Neurol 56, 598–609. 10.1016/0014-4886(77)90323-5.

54. Dembrow, N.C., Sawchuk, S., Dalley, R., Opitz-Araya, X., Hudson, M., Radaelli, C., Alfiler, L., Walling-Bell, S., Bertagnolli, D., Goldy, J., et al. (2024). Areal specializations in the morpho-electric and transcriptomic properties of primate layer 5 extratelencephalic projection neurons. Cell Reports 43. 10.1016/j.celrep.2024.114718.

55. Jacobs, B., Garcia, M.E., Shea-Shumsky, N.B., Tennison, M.E., Schall, M., Saviano, M.S., Tummino, T.A., Bull, A.J., Driscoll, L.L., Raghanti, M.A., et al. (2017). Comparative morphology of gigantopyramidal neurons in primary motor cortex across mammals. The Journal of comparative neurology 526, 496–536. 10.1002/cne.24349.

56. Kaiserman-Abramof, I.R., and Peters, A. (1972). Some aspects of the morphology of Betz cells in the cerebral cortex of the cat. Brain Research 43, 527–546. 10.1016/0006-8993(72)90406-4.

57. Fuentealba-Villarroel, F.J., Renner, J., Hilbig, A., Bruton, O.J., and Rasia-Filho, A.A. (2022). Spindle-Shaped Neurons in the Human Posteromedial (Precuneus) Cortex. Front. Synaptic Neurosci. 13. 10.3389/fnsyn.2021.769228.

58. Suarez-Sola, M.L., GonzalezDelgado, F.J., Pueyo-Morlans, M., Medina-Bolivar, C., HernandezAcosta, N.C., Gonzalez-Gomez, M., and Meyer, G. (2009). Neurons in the white matter of the adult human neocortex. Front. Neuroanat. 3. 10.3389/neuro.05.007.2009.

59. Zolnik, T.A., Bronec, A., Ross, A., Staab, M., Sachdev, R.N.S., Molnár, Z., Eickholt, B.J., and Larkum, M.E. (2024). Layer 6b controls brain state via apical dendrites and the higher-order thalamocortical system. Neuron 112, 805–820.e4. 10.1016/j.neuron.2023.11.021.

60. Radaelli, C., Schmitz, M., Liu, X.-P., Sawchuk, S., Opitz-Araya, X., Hudson, M., Taskin, N., Bertagnolli, D., Goldy, J., Ko, A.L., et al. (2025). HCN channels reveal conserved and divergent physiology in supragranular pyramidal neurons in primate species. Preprint at bioRxiv, 10.1101/2025.08.22.671856 https://doi.org/10.1101/2025.08.22.671856.

61. Mohan, H., Verhoog, M.B., Doreswamy, K.K., Eyal, G., Aardse, R., Lodder, B.N., Goriounova, N.A., Asamoah, B., B. Brakspear, A.B.C., Groot, C, et al. (2015). Dendritic and Axonal Architecture of Individual Pyramidal Neurons across Layers of Adult Human Neocortex. Cerebral Cortex 25, 4839–4853. 10.1093/cercor/bhv188.

62. Moradi Chameh, H., Rich, S., Wang, L., Chen, F.-D., Zhang, L., Carlen, P.L., Tripathy, S.J., and Valiante, T.A. (2021). Diversity amongst human cortical pyramidal neurons revealed via their sag currents and frequency preferences. Nat Commun 12, 2497. 10.1038/s41467-021-22741-9.

63. Benavides-Piccione, R., Blazquez-Llorca, L., Kastanauskaite, A., Fernaud-Espinosa, I., Tapia-González, S., and DeFelipe, J. (2024). Key morphological features of human pyramidal neurons. Cereb Cortex 34, bhae180. 10.1093/cercor/bhae180.

64. Gilman, J.P., Medalla, M., and Luebke, J.I. (2017). Area-Specific Features of Pyramidal Neurons-a Comparative Study in Mouse and Rhesus Monkey. Cereb Cortex 27, 2078–2094. 10.1093/cercor/bhw062.

65. Gao, L., Liu, S., Gou, L., Hu, Y., Liu, Y., Deng, L., Ma, D., Wang, H., Yang, Q., Chen, Z., et al. (2022). Single-neuron projectome of mouse prefrontal cortex. Nat Neurosci 25, 515–529. 10.1038/s41593-022-01041-5.

66. Muñoz-Castañeda, R., Zingg, B., Matho, K.S., Chen, X., Wang, Q., Foster, N.N., Li, A., Narasimhan, A., Hirokawa, K.E., Huo, B., et al. (2021). Cellular anatomy of the mouse primary motor cortex. Nature 598, 159–166. 10.1038/s41586-021-03970-w.

67. Hutsler, J.J., Lee, D.-G., and Porter, K.K. (2005). Comparative analysis of cortical layering and supragranular layer enlargement in rodent carnivore and primate species. Brain Research 1052, 71–81. 10.1016/j.brainres.2005.06.015.

68. Hofman, M.A. (1988). Size and shape of the cerebral cortex in mammals. II. The cortical volume. Brain Behav Evol 32, 17–26. 10.1159/000116529.

69. Driessens, S.L.W., Galakhova, A.A., Heyer, D.B., Pieterse, I.J., Wilbers, R., Mertens, E.J., Waleboer, F., Heistek, T.S., Coenen, L., Meijer, J.R., et al. (2023). Genes associated with cognitive ability and HAR show overlapping expression patterns in human cortical neuron types. Nat Commun 14, 4188. 10.1038/s41467-023-39946-9.

70. Starr, A.L., and Fraser, H.B. (2025). A General Principle of Neuronal Evolution Reveals a Human-Accelerated Neuron Type Potentially Underlying the High Prevalence of Autism in Humans. Mol Biol Evol 42, msaf189. 10.1093/molbev/msaf189.

71. Heyer, D.B., Wilbers, R., Galakhova, A.A., Hartsema, E., Braak, S., Hunt, S., Verhoog, M.B., Muijtjens, M.L., Mertens, E.J., Idema, S., et al. (2022). Verbal and General IQ Associate with Supragranular Layer Thickness and Cell Properties of the Left Temporal Cortex. Cereb Cortex 32, 2343–2357. 10.1093/cercor/bhab330.

72. Won, H., Huang, J., Opland, C.K., Hartl, C.L., and Geschwind, D.H. (2019). Human evolved regulatory elements modulate genes involved in cortical expansion and neurodevelopmental disease susceptibility. Nat Commun 10, 2396. 10.1038/s41467-019-10248-3.

73. Pressl, C., Mätlik, K., Kus, L., Darnell, P., Luo, J.-D., Paul, M.R., Weiss, A.R., Liguore, W., Carroll, T.S., Davis, D.A., et al. (2024). Selective vulnerability of layer 5a corticostriatal neurons in Huntington’s disease. Neuron 112, 924–941.e10. 10.1016/j.neuron.2023.12.009.

74. Dick, F., Johanson, G.A.S., Tysnes, O.-B., Alves, G., Dölle, C., and Tzoulis, C. (2025). Brain Proteome Profiling Reveals Common and Divergent Signatures in Parkinson’s Disease, Multiple System Atrophy, and Progressive Supranuclear Palsy. Mol Neurobiol 62, 2801–2816. 10.1007/s12035-024-04422-y.

75. Marcassa, G., Dascenco, D., Lorente-Echeverría, B., Daaboul, D., Vandensteen, J., Leysen, E., Baltussen, L., Howden, A.J.M., and de Wit, J. (2025). Synaptic signatures and disease vulnerabilities of layer 5 pyramidal neurons. Nat Commun 16, 228. 10.1038/s41467-024-55470-w.

76. Goralski, T.M., Meyerdirk, L., Breton, L., Brasseur, L., Kurgat, K., DeWeerd, D., Turner, L., Becker, K., Adams, M., Newhouse, D.J., et al. (2024). Spatial transcriptomics reveals molecular dysfunction associated with cortical Lewy pathology. Nat Commun 15, 2642. 10.1038/s41467-024-47027-8.

77. Kumar, P., and Ohana, O. (2008). Inter- and Intralaminar Subcircuits of Excitatory and Inhibitory Neurons in Layer 6a of the Rat Barrel Cortex. Journal of Neurophysiology 100, 1909–1922. 10.1152/jn.90684.2008.

78. Huang, S., Wu, S.J., Sansone, G., Ibrahim, L.A., and Fishell, G. (2024). Layer 1 neocortex: Gating and integrating multidimensional signals. Neuron 112, 184–200. 10.1016/j.neuron.2023.09.041.

79. Jones, K.E., Angielczyk, K.D., and Pierce, S.E. (2019). Stepwise shifts underlie evolutionary trends in morphological complexity of the mammalian vertebral column. Nat Commun 10, 5071. 10.1038/s41467-019-13026-3.

80. Gast, R., Solla, S.A., and Kennedy, A. (2024). Neural heterogeneity controls computations in spiking neural networks. Proc Natl Acad Sci U S A 121, e2311885121. 10.1073/pnas.2311885121.

81. Ting, J.T., Daigle, T.L., Chen, Q., and Feng, G. (2014). Acute Brain Slice Methods for Adult and Aging Animals: Application of Targeted Patch Clamp Analysis and Optogenetics. In Patch-Clamp Methods and Protocols Methods in Molecular Biology., M. Martina and S. Taverna, eds. (Springer), pp. 221–242. 10.1007/978-1-4939-1096-0_14.

82. Neher, E., and Sakmann, B. (1992). The patch clamp technique. Sci Am 266, 44–51. 10.1038/scientificamerican0392-44.

83. Fonov, V.S., Evans, A.C., McKinstry, R.C., Almli, C.R., and Collins, D.L. (2009). Unbiased nonlinear average age-appropriate brain templates from birth to adulthood. NeuroImage, S102.

84. Ding, S.-L., Royall, J.J., Sunkin, S.M., Ng, L., Facer, B.A.C., Lesnar, P., Guillozet-Bongaarts, A., McMurray, B., Szafer, A., Dolbeare, T.A., et al. (2016). Comprehensive cellular-resolution atlas of the adult human brain. Journal of Comparative Neurology 524, 3127–3481. 10.1002/cne.24080.

85. Peng, H., Ruan, Z., Long, F., Simpson, J.H., and Myers, E.W. (2010). V3D enables real-time 3D visualization and quantitative analysis of large-scale biological image data sets. Nat Biotechnol 28, 348–353. 10.1038/nbt.1612.

86. Zhou, Z., Liu, X., Long, B., and Peng, H. (2016). TReMAP: Automatic 3D Neuron Reconstruction Based on Tracing, Reverse Mapping and Assembling of 2D Projections. Neuroinform 14, 41–50. 10.1007/s12021-015-9278-1.

87. Gliko, O., Mallory, M., Dalley, R., Gala, R., Gornet, J., Zeng, H., Sorensen, S.A., and Sümbül, U. (2024). High-throughput analysis of dendrite and axonal arbors reveals transcriptomic correlates of neuroanatomy. Nat Commun 15, 6337. 10.1038/s41467-024-50728-9.

88. Peng, H., Bria, A., Zhou, Z., Iannello, G., and Long, F. (2014). Extensible visualization and analysis for multidimensional images using Vaa3D. Nat Protoc 9, 193–208. 10.1038/nprot.2014.011.

89. Bria, A., Iannello, G., Onofri, L., and Peng, H. (2016). TeraFly: real-time three-dimensional visualization and annotation of terabytes of multidimensional volumetric images. Nat Methods 13, 192–194. 10.1038/nmeth.3767.

90. Hao, Y., Hao, S., Andersen-Nissen, E., Mauck, W.M., Zheng, S., Butler, A., Lee, M.J., Wilk, A.J., Darby, C., Zager, M., et al. (2021). Integrated analysis of multimodal single-cell data. Cell 184, 3573–3587.e29. 10.1016/j.cell.2021.04.048.

91. Becht, E., McInnes, L., Healy, J., Dutertre, C.A., Kwok, I.W.H., Ng, L.G., Ginhoux, F., and Newell, E.W. (2019). Dimensionality reduction for visualizing single-cell data using UMAP. Nature Biotechnology 37, 38–47. 10.1038/nbt.4314.

92. Chitra, U., Arnold, B.J., Sarkar, H., Sanno, K., Ma, C., Lopez-Darwin, S., and Raphael, B.J. (2025). Mapping the topography of spatial gene expression with interpretable deep learning. Nat Methods 22, 298–309. 10.1038/s41592-024-02503-3.

93. Egger, V., Nevian, T., and Bruno, R.M. (2008). Subcolumnar Dendritic and Axonal Organization of Spiny Stellate and Star Pyramid Neurons within a Barrel in Rat Somatosensory Cortex. Cerebral Cortex 18, 876–889. 10.1093/cercor/bhm126.

94. Scorcioni, R., Polavaram, S., and Ascoli, G.A. (2008). L-Measure: a web-accessible tool for the analysis, comparison and search of digital reconstructions of neuronal morphologies. Nat Protoc 3, 866–876. 10.1038/nprot.2008.51.

95. Markram, H., Muller, E., Ramaswamy, S., Reimann, M.W., Abdellah, M., Sanchez, C.A., Ailamaki, A., Alonso-Nanclares, L., Antille, N., Arsever, S., et al. (2015). Reconstruction and Simulation of Neocortical Microcircuitry. Cell 163, 456–492. 10.1016/j.cell.2015.09.029.

96. Traag, V.A., Waltman, L., and van Eck, N.J. (2019). From Louvain to Leiden: guaranteeing well-connected communities. Sci Rep 9, 5233. 10.1038/s41598-019-41695-z.

