## Supplemental Figures for "Human Neocortical Glutamatergic Neurons Revealed Through Multimodal Profiling"

### Figure S1 Transcriptomic quality control and cortical lobe distribution

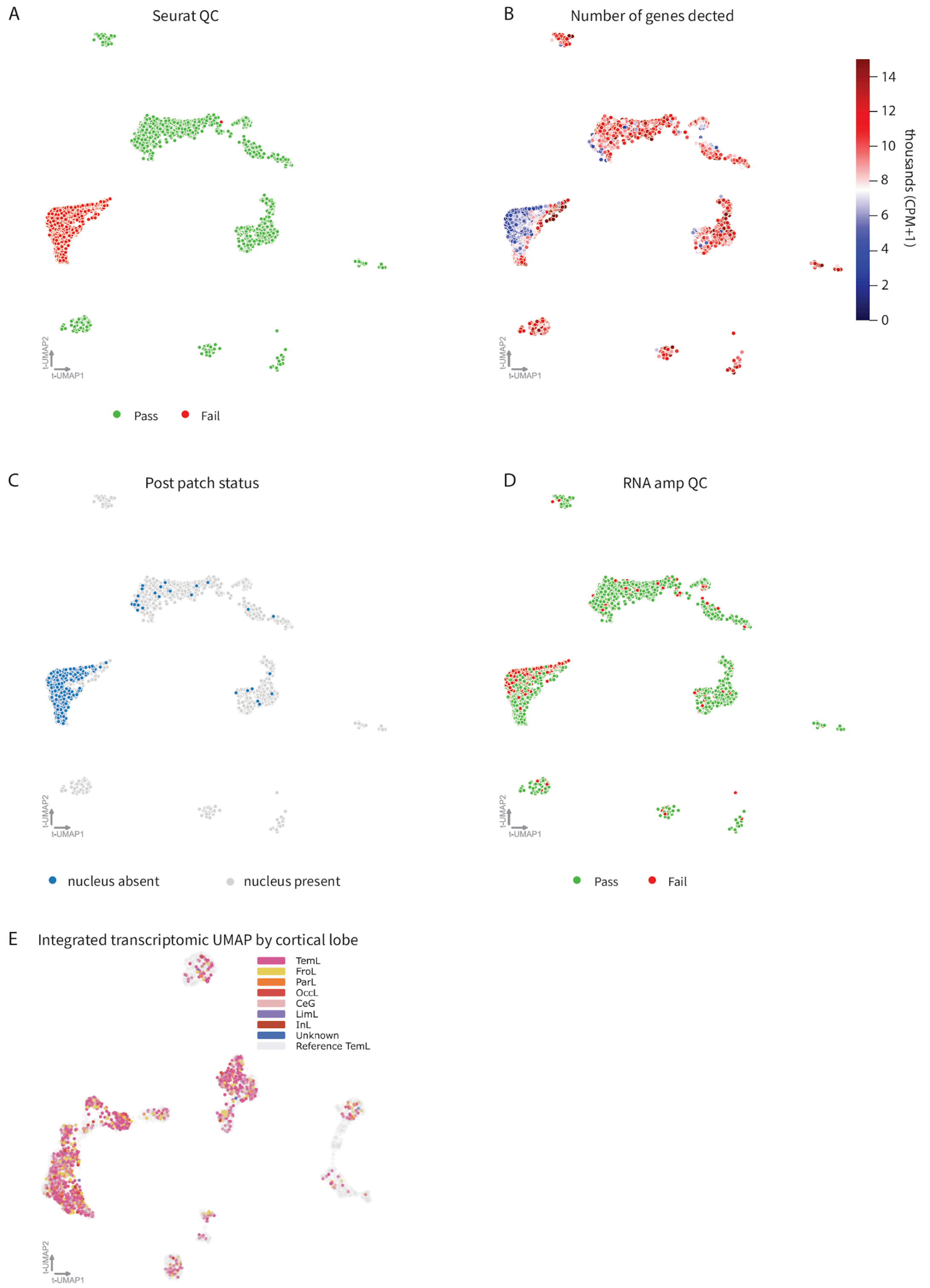


**Figure S1** Transcriptomic quality control and cortical lobe distribution

Integrated UMAP of reference and Patch-seq transcriptomes before removal of misaligned Patch-seq neurons, color-coded by Seurat QC (A), number of genes detected (B), post-Patch status (C), and RNA-amplification QC (D). All metrics indicate an island of low-quality cells that were removed for the study (Seurat QC = Fail). (E) Integrated UMAP of reference and Patch-seq transcriptomes after removal of misaligned Patch-seq neurons, as seen in Figure 1C, colored by cortical lobe made with Cytosplore Viewer.

### Figure S2 Patch-seq morphology reconstructions

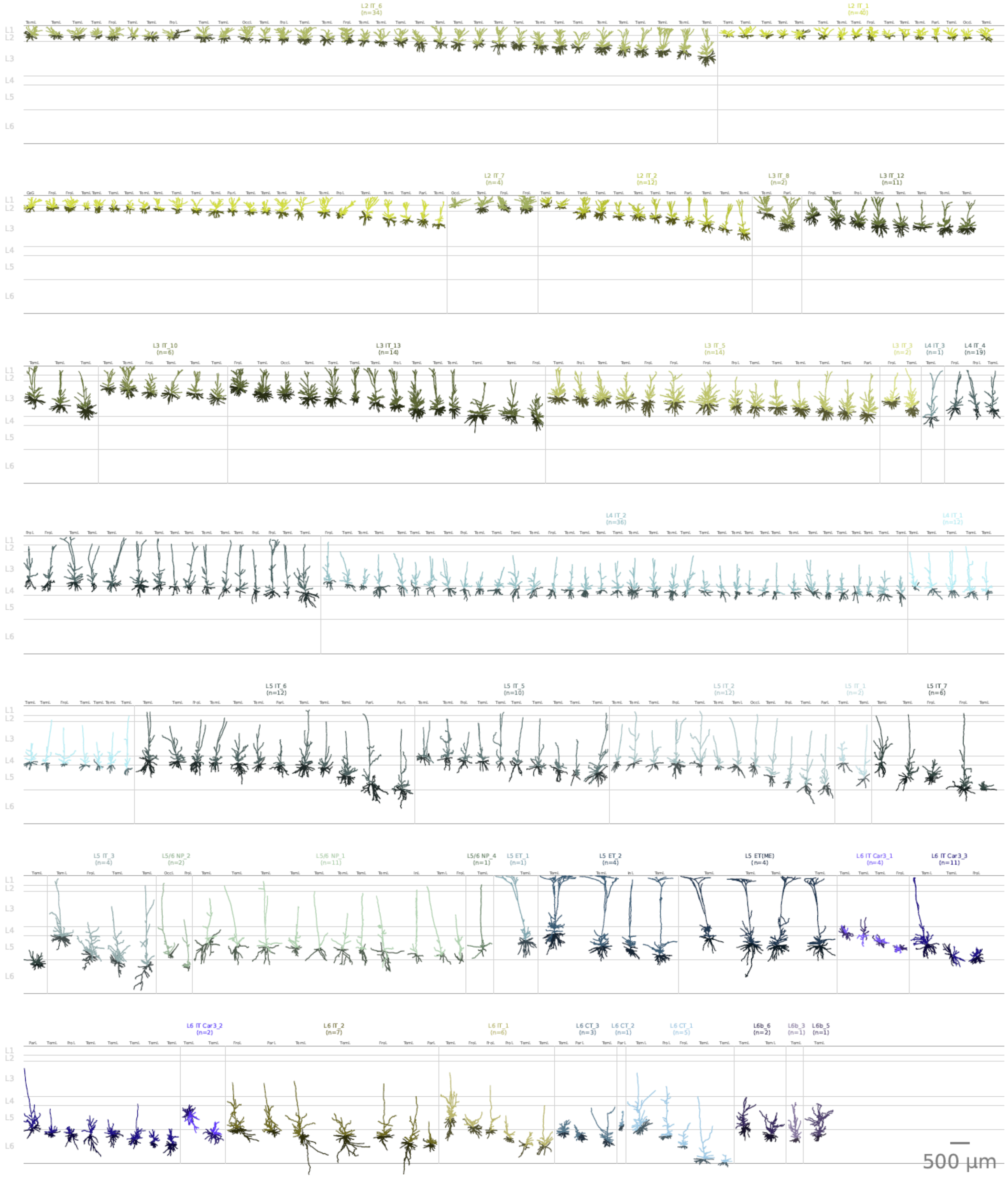


### Figure S2 Patch-seq morphology reconstructions

Reconstructions showing apical dendrites in supertypes colors and basal dendrites in a darker variant of the same color aligned to an average layer template.

#### Figure S2 continued Patch-seq morphology reconstructions

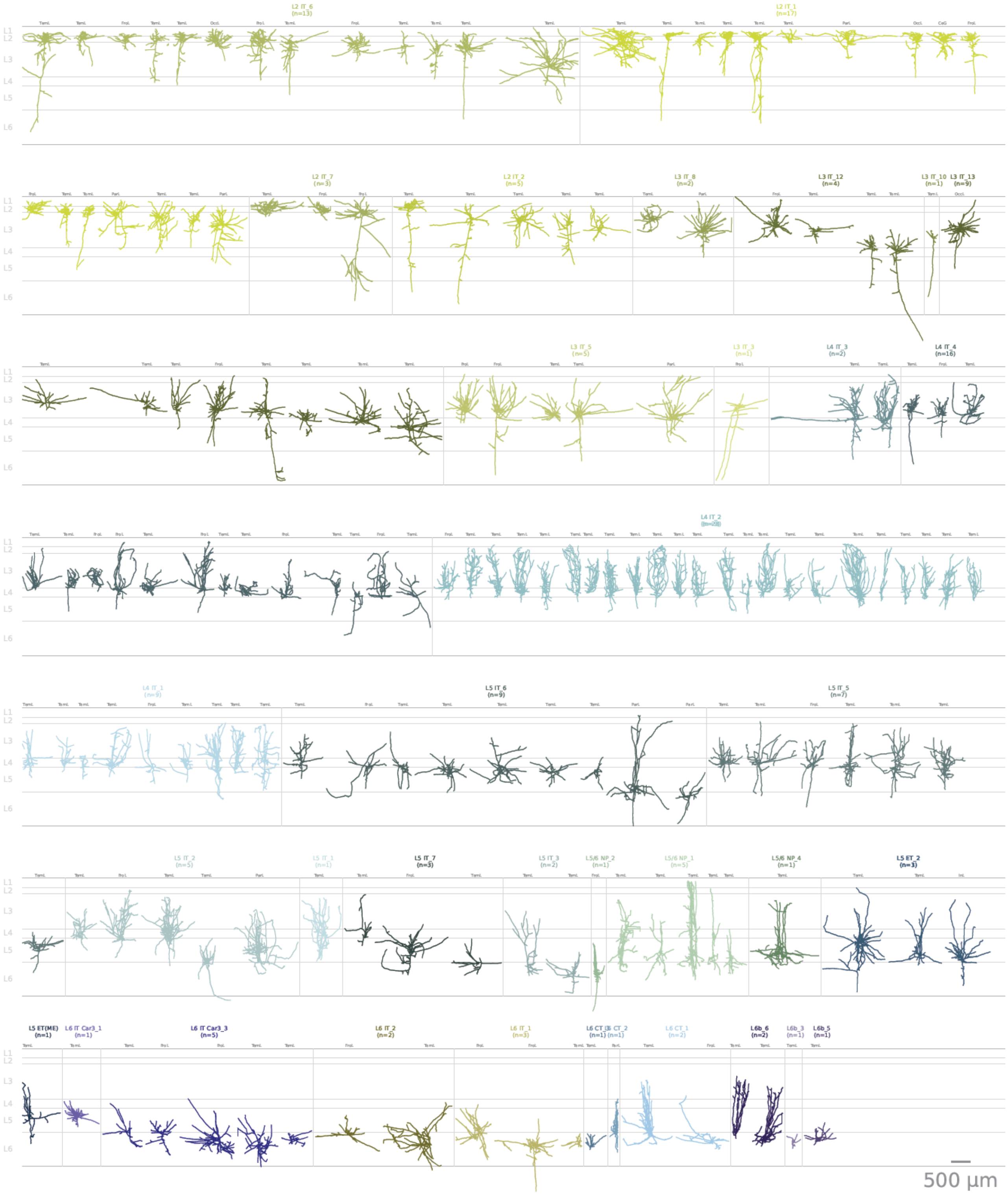


#### Figure S2 continued Patch-seq morphology reconstructions

Axon reconstructions colored by supertype identity aligned to an average layer template.

### Figure S3 Neurons recorded in acute and slice culture paradigms

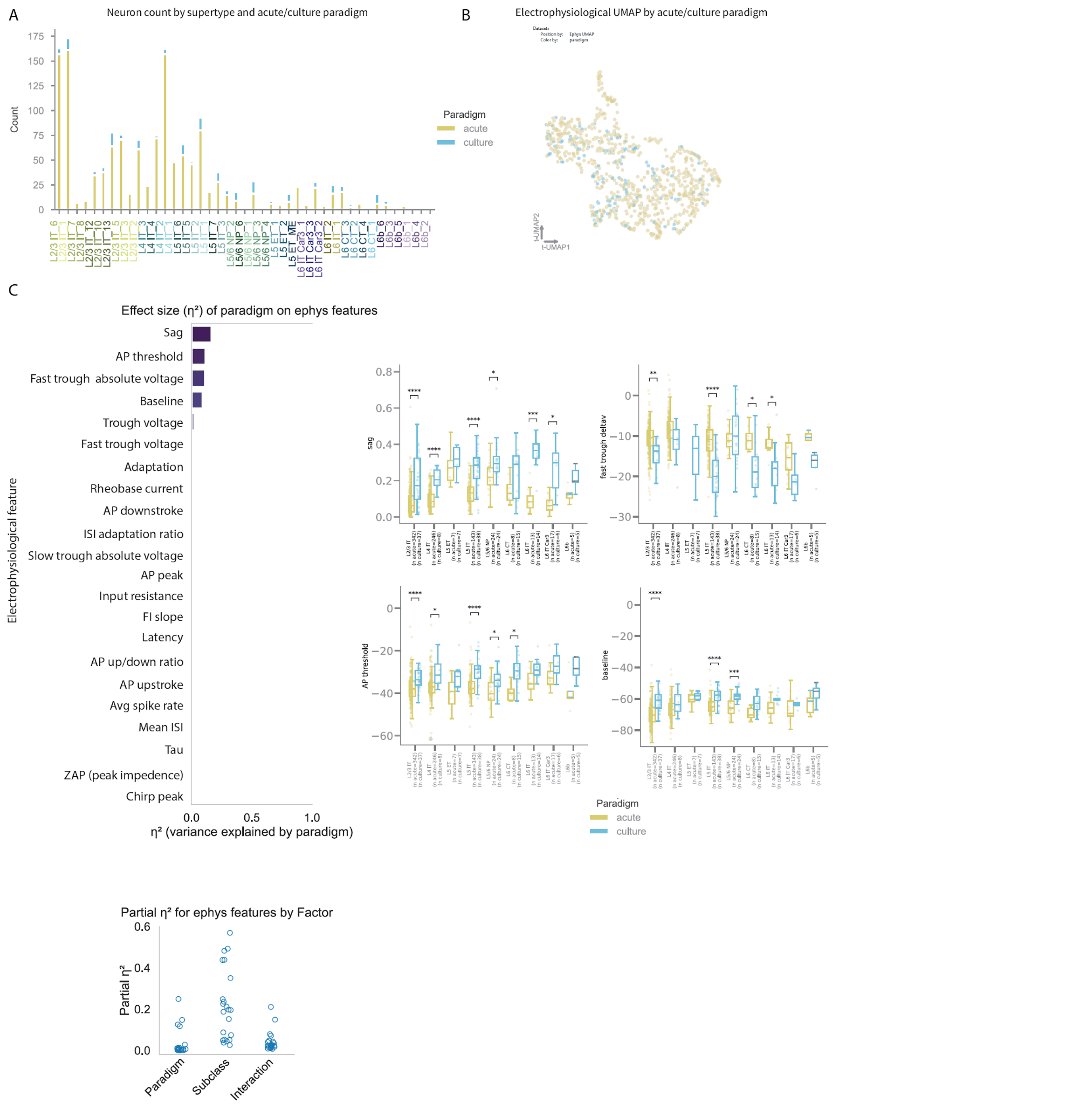


### Figure S3 Neurons recorded in acute and slice culture paradigms

A. Neuron count across supertype by acute/culture paradigm.

B. Electrophysiological UMAP colored by acute/culture paradigm.

C. Two-way ANOVA of electrophysiological features examining the effect of paradigm. Bars represent the proportion of variance explained by paradigm (η²), sorted in descending order. Distributions of partial η² values for features across paradigm, subclass and interaction are shown. Box plots depict the top four features most affected by paradigm, showing their distributions across subclasses. Statistical details are provided in Table S2.

### Figure S4 Subclass analysis excluding soma cortical depth and matched-neuron classification

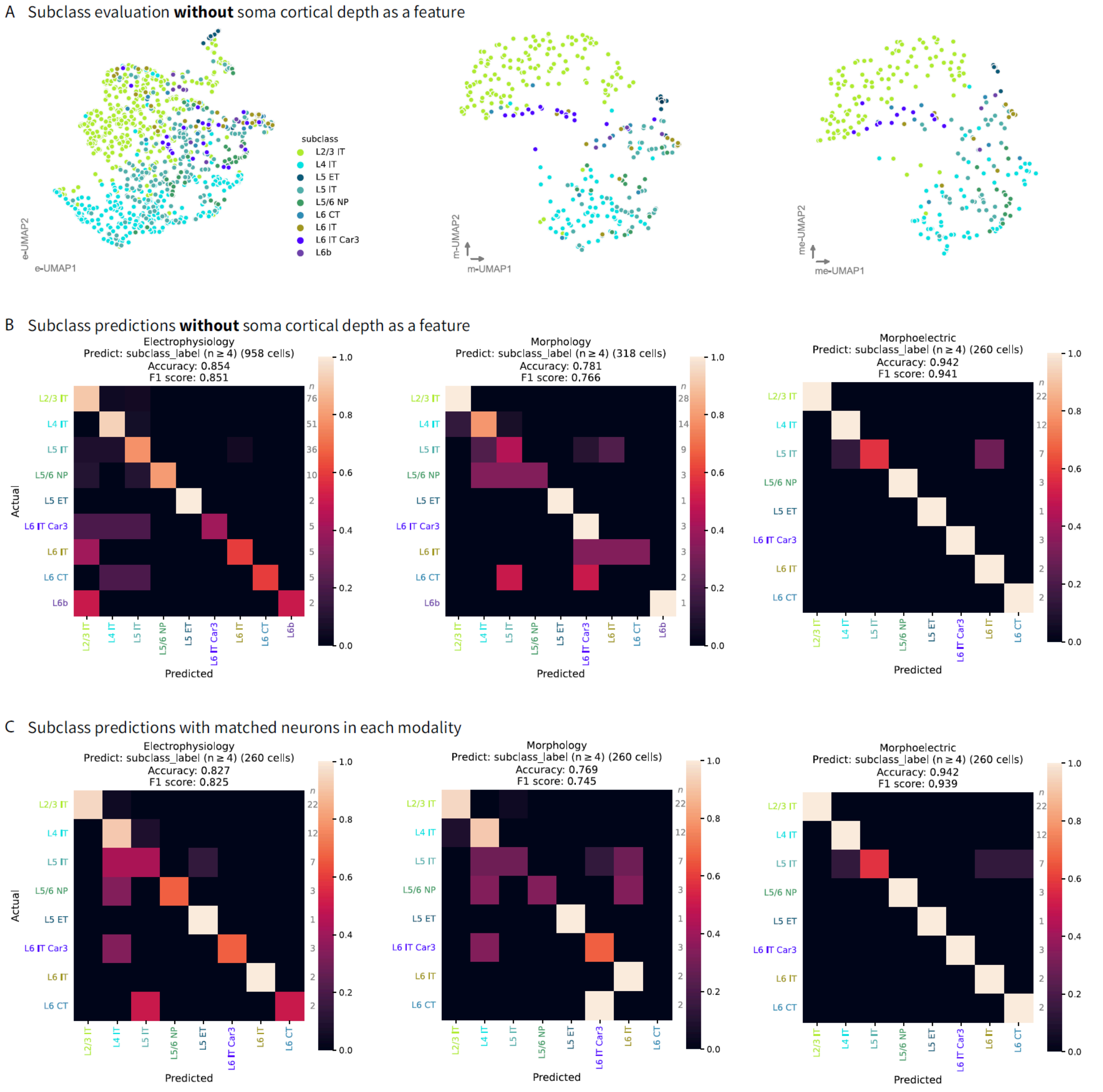


Figure S4 Subclass analysis excluding soma cortical depth and matched-neuron classification

A. UMAP representations of electrophysiological (E), morphological (M), and combined morphoelectric (ME) features with soma cortical depth features removed.

B. Confusion matrix of a random forest classifier predicting subclass from E, M, and combined ME features with soma cortical depth features removed.

C. Confusion matrix of a random forest classifier predicting subclass from E, M, and combined ME features using a matched set of neurons.

### Figure S5 Additional L2 and L3 morphoelectric analysis

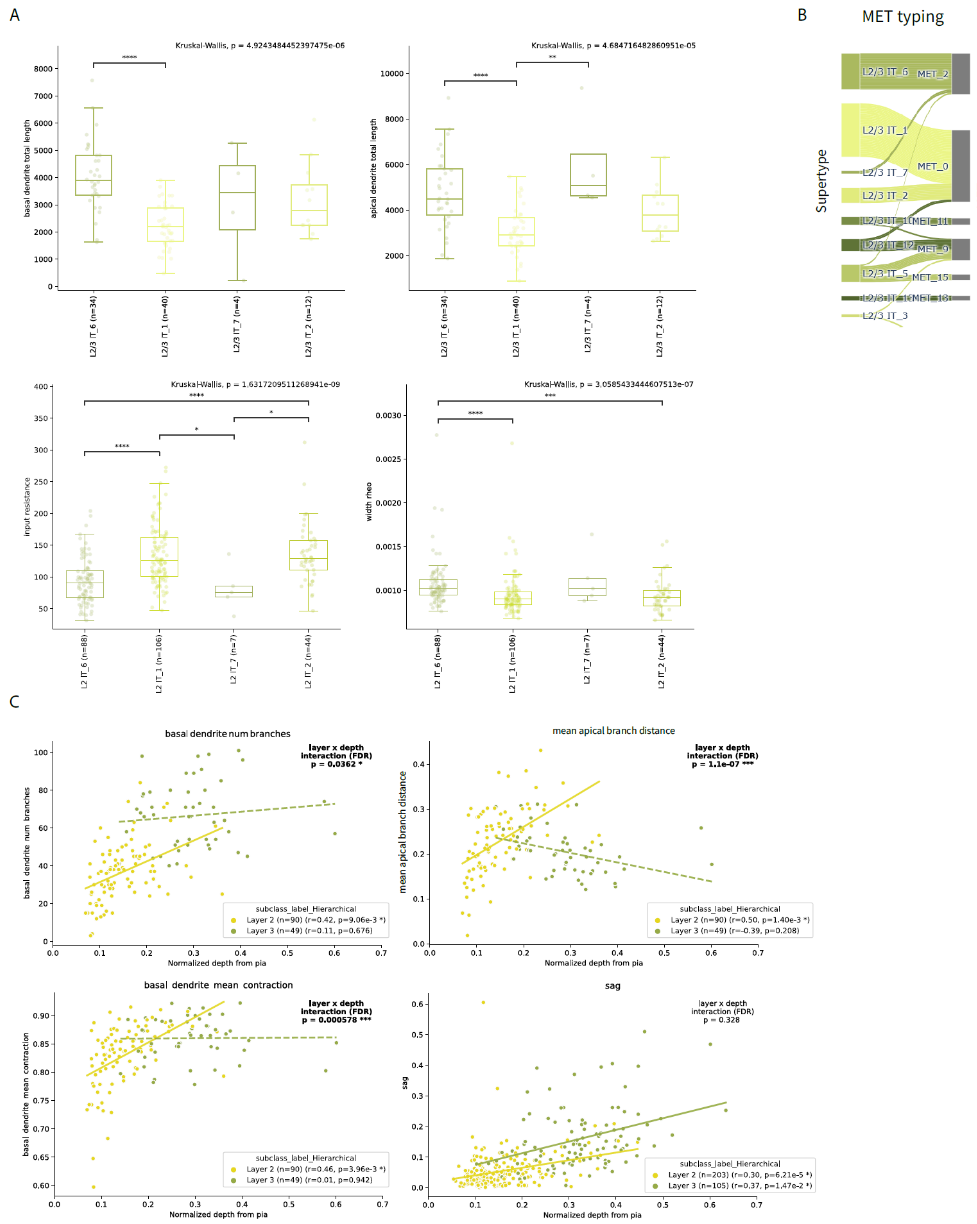


### Figure S5 Additional L2 and L3 morphoelectric analysis

A. Morphological (top) human L2 supertypes with morphoelectric properties with the same relative relationship as in mouse; electrophysiological (bottom) properties that distinguish L2 supertypes. Statistical comparisons of features across supertypes were performed using the Kruskal-Wallis H-test, with FDR correction. For features with significant Kruskal-Wallis results ($p < 0.05$, FDR-corrected), post-hoc pairwise comparisons were conducted using Dunn's test with FDR correction. Full statistical details are provided in Table S2**.**

B. River plot showing the relationships between L2 and L3 supertypes (left) and assigned MET-types (right) for neurons from Patch-seq recordings with all three data modalities available.

C. Scatter plot of morphological and electrophysiological features by cortical depth, color-coded by layer. Two-way ANOVA tested effects of depth from pia, cortical layer, and their interaction; significant ($p<0.05$) FDR (Benjamini–Hochberg procedure) corrected depth-layer interactions are shown in bold in the upper right. For each layer, a linear regression (ordinary least squares) with HC3 heteroscedasticity-robust standard errors was fit to relate the feature to depth. The corresponding fitted line is plotted for each layer, shown as solid when the robust F-test P value is significant ($p<0.05$) after FDR correction (Benjamini–Hochberg procedure) and dashed when it is not. The Pearson correlation (r) and FDR-corrected P value for each layer are reported in the legend. Full statistical details are provided in Table S2**.**

#### Figure S6 Alignment of prior taxonomy and published Patch-seq data with the SEA-AD taxonomy

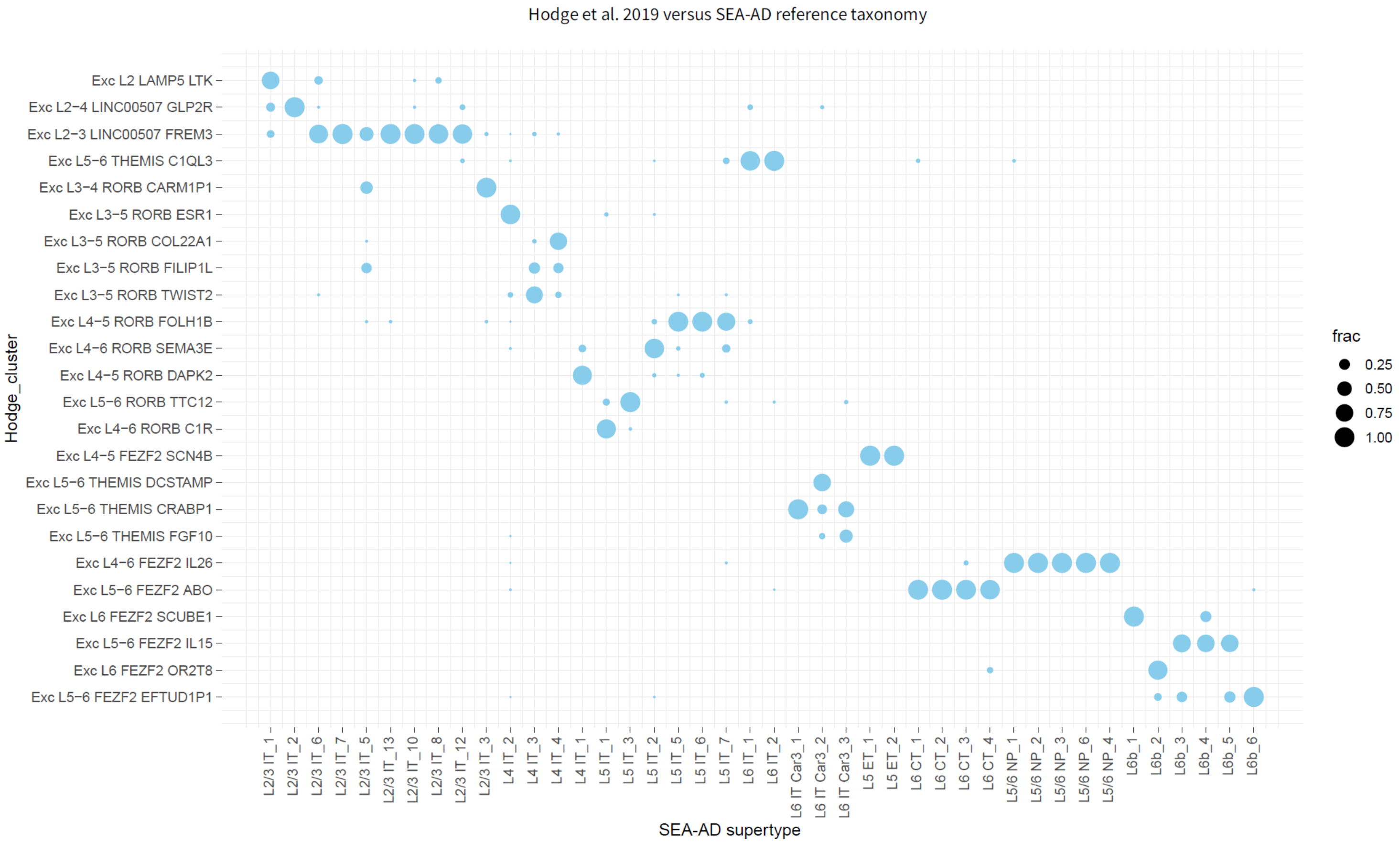


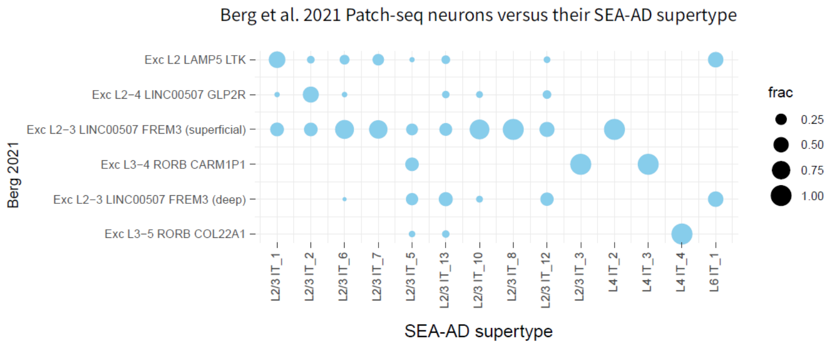


#### Figure S6 Alignment of prior taxonomy and published Patch-seq data with the SEA-AD taxonomy

Hodge et al. 2019 vs SEA-AD taxonomy, Berg et al. 2021 Patch-seq neurons to SEA-AD taxonomy

#### Figure S7 Layer 2 and 3 classifiers to predict supertype and d prime

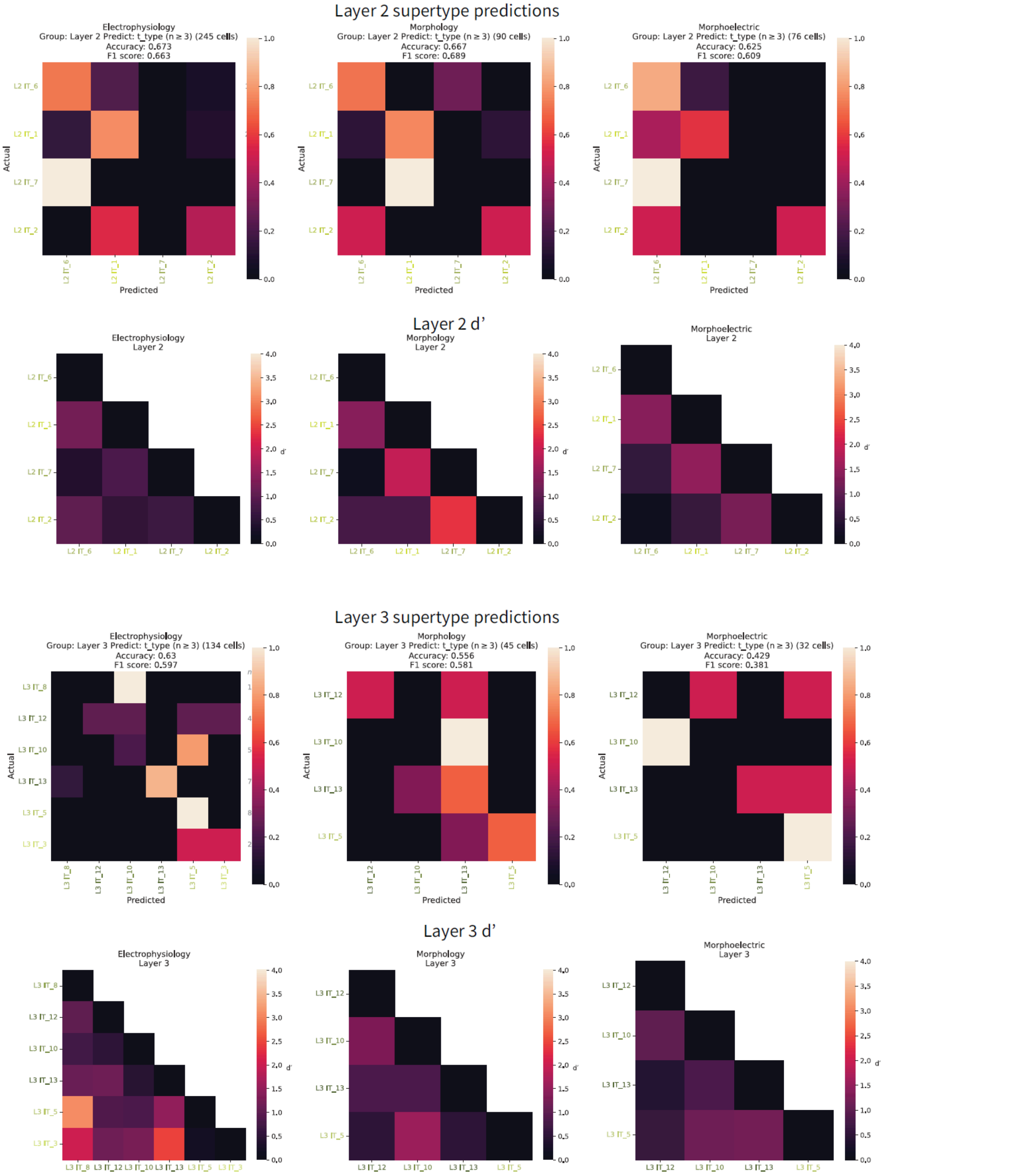


#### Figure S7 Layer 2 and 3 classifiers to predict supertype and d prime

1st and 3rd row - Random Forest classifier to predict L2 and L3 supertypes, respectively.

2nd and 4th row - Pairwise L2 and L3 supertype, respectively, comparisons using the d' discriminability metric, where higher values indicate greater separation.

### Figure S8 Additional L4 morphoelectric analysis

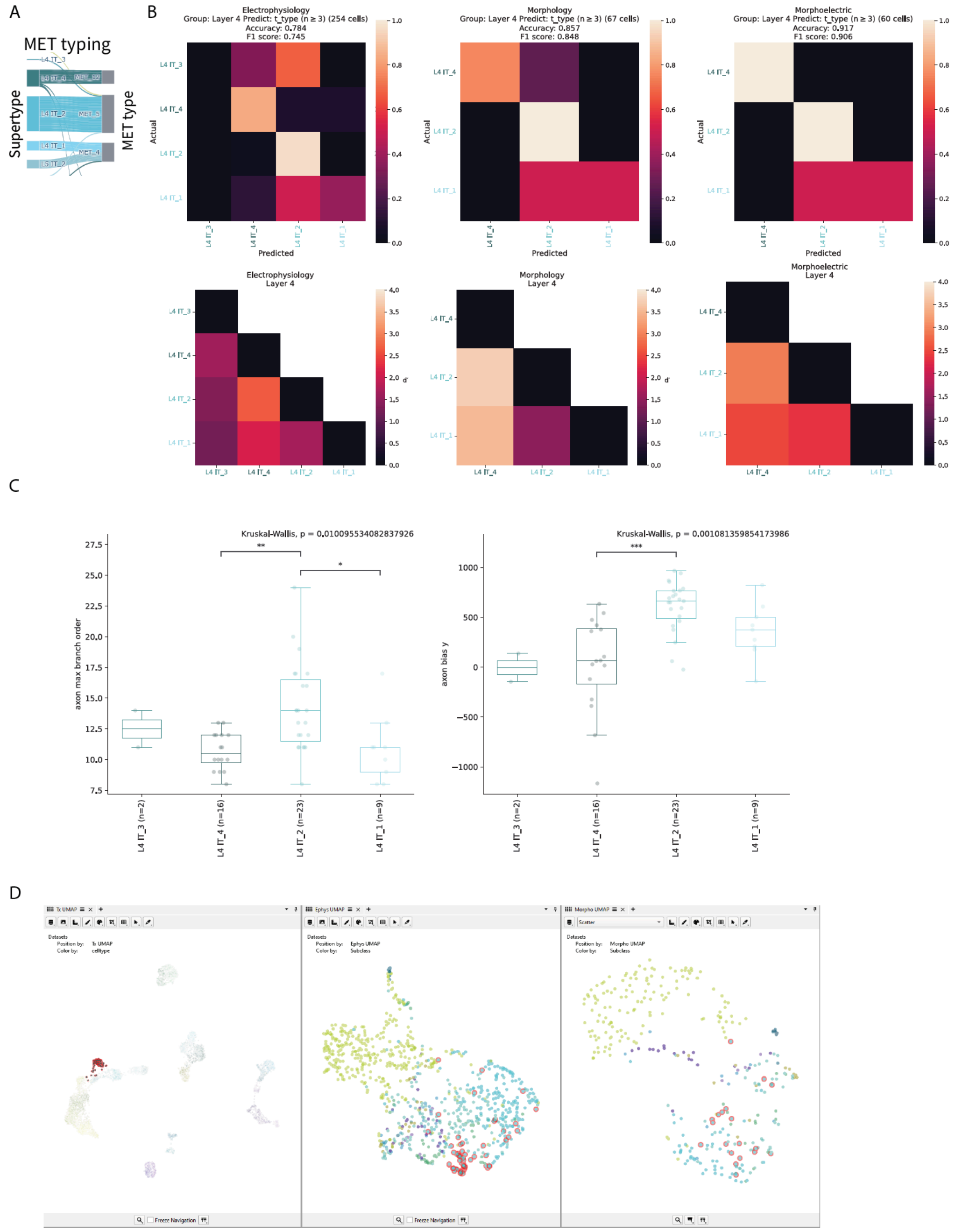


### Figure S8 Additional L4 morphoelectric analysis

A. River plot showing the relationships between L4 supertypes (left) and assigned MET-types (right) for neurons from Patch-seq recordings with all three data modalities available.

B. Top row - Random Forest classifier to predict L4 IT supertypes, Bottom row- Pairwise L4 IT supertype comparisons using the d′ discriminability metric, where higher values indicate greater separation.

C. Axon features, maximum branch order and vertical bias, highlighted the differences across L4 supertypes. Kruskal-Wallis H-test on morphology axon features between supertypes in the same layer. FDR corrected. If KW FDR is significant ($p < 0.05$) post-hoc Dunn test on pairwise supertypes within layer. FDR corrected. Full statistical details are provided in Table S2**.**

D. A screen capture from the Cytosplore Viewer highlighting L4 IT_4.

### Figure S9 Additional L5 morphoelectric analysis

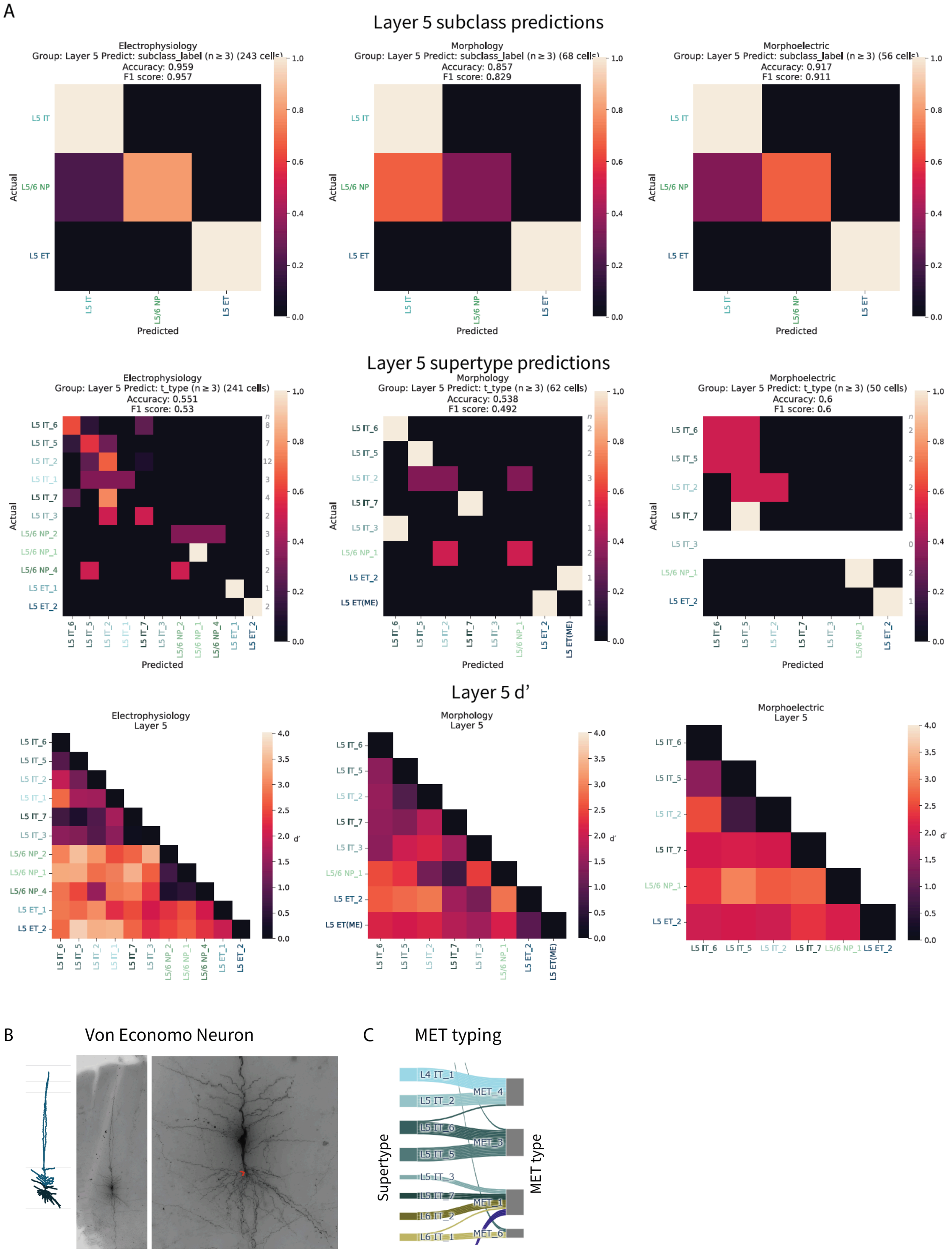


### Figure S9 Additional L5 morphoelectric analysis

A. *Top and middle row* - Random Forest classifier to predict L5 subclasses and supertypes respectively, *bottom row*- Pairwise L5 supertype comparisons using the d′ discriminability metric, where higher values indicate greater separation.
B. Reconstruction of a Von Economo neuron (left), with high resolution overview (middle) and a zoom in of the perisomatic region showing the axon origin (red arrow).

C. River plot showing the relationships between L5 supertypes (left) and assigned MET-types (right) for neurons from Patch-seq recordings with all three data modalities available.

### Figure S10 Additional L6 morphoelectric analysis

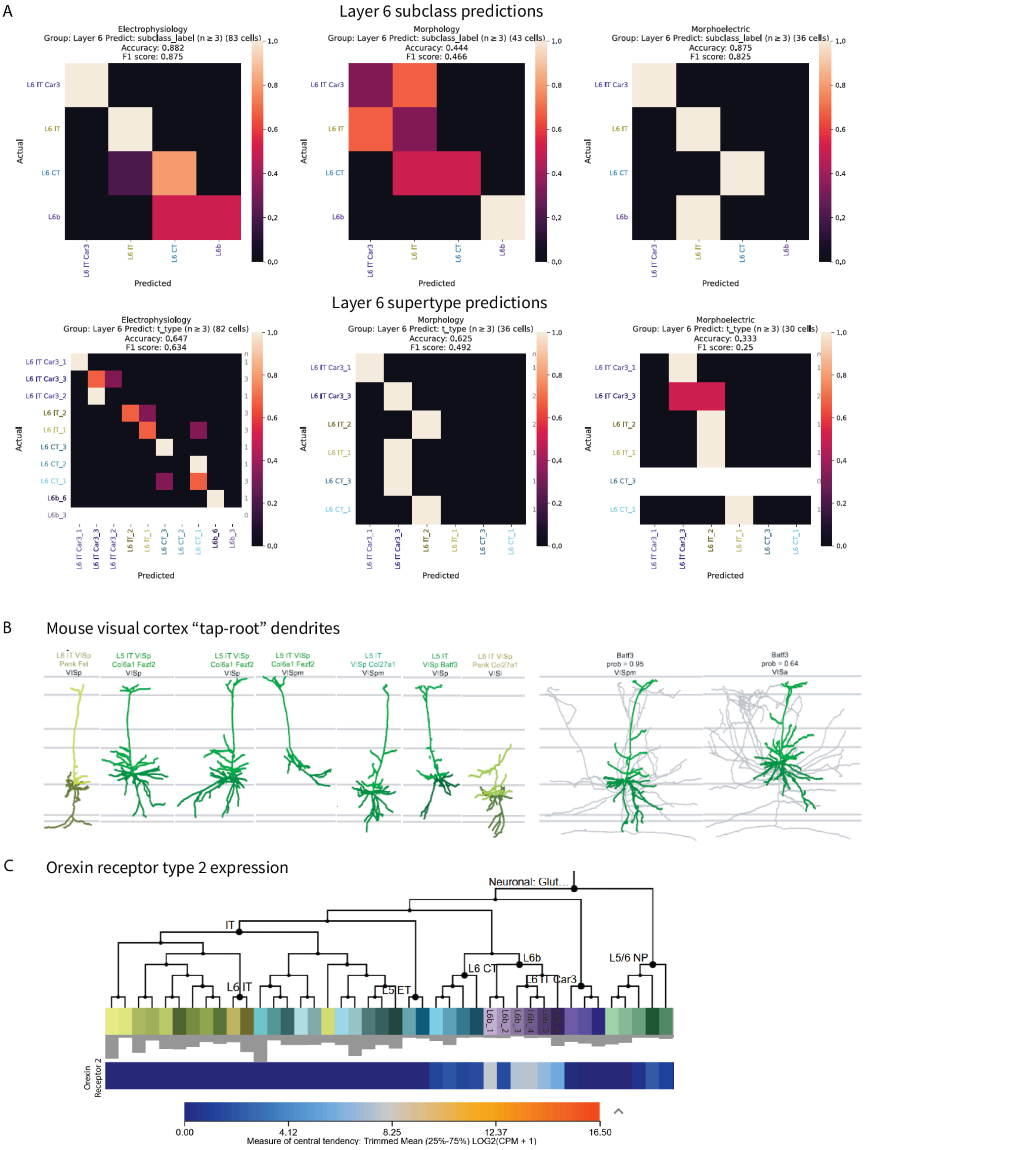


### Figure S10 Additional L6 morphoelectric analysis

A. Top row - Random Forest classifier to predict L6 IT supertypes, Bottom row- Pairwise L6 IT supertype comparisons using the d′ discriminability metric, where higher values indicate greater separation.
B. Mouse visual cortex neurons with “tap-root” dendrites modified from Sorensen et al. (manuscript in revision)
C. Orexin receptor type 2 expression from SEA-AD snRNA-seq reference data (https://celltypes.brain-map.org/rnaseq/Human-MTG-10x_SEA-AD).

### Figure S11 Additional cross-species morphoelectric analysis

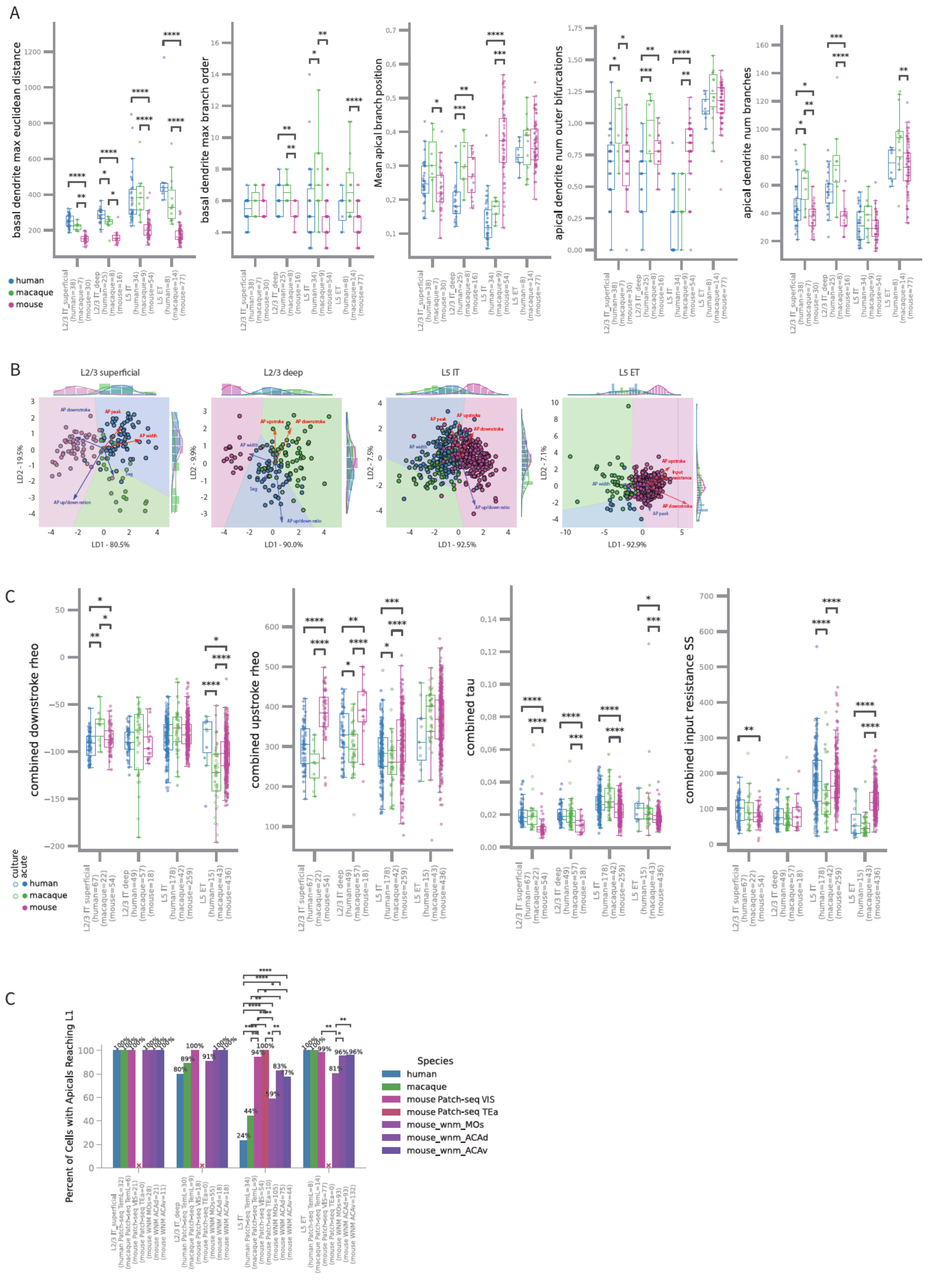


### Figure S11 Additional cross-species morphoelectric analysis

A. Morphological feature comparison across species and subclass. Mann-Whitney U-test when only two species groups, otherwise Kruskal-Wallis H-test, FDR corrected. If KW FDR was significant ($p < 0.05$) post-hoc Dunn test on pairwise species groups, FDR corrected. Full statistical details are provided in Table S2.

B. Linear Discriminant Analysis (LDA) of species-specific electrophysiological features in L2/3 superficial and deep, L5 IT and L5 ET subclasses. The first two linear discriminates and their explained variance are plotted on the x- and y-axis. Transparent colored regions represent decision boundaries derived from the LDA model and the five features with the largest combined contributions are overlaid as arrows. Direction, length and color (red: positive, blue: negative) of arrow reflects relative contribution of each feature.
C. Electrophysiological feature comparison with the top electrophysiological features most affected by species depicted, culture denoted as open circles and acute as closed circles. Mann-Whitney U-test when only two species groups, otherwise Kruskal-Wallis H-test, FDR corrected. If KW FDR is significant ($p < 0.05$) post-hoc Dunn test on pairwise species groups, FDR corrected. Full statistical details are provided in Table S2.
D. Bar plot with the percentage of neurons with apicals reaching L1 across species with significance determined with Chi-square test of independence.
